# Cell cycle-dependent active stress drives epithelia remodeling

**DOI:** 10.1101/804294

**Authors:** John Devany, Daniel M. Sussman, Takaki Yamamoto, M. Lisa Manning, Margaret L. Gardel

**Affiliations:** Department of Physics, James Franck Institute and Institute for Biophysical Dynamics, The University of Chicago, Chicago, IL 60637, USA; Department of Physics and BioInspired Institute, Syracuse University, Syracuse, NY 13244, USA; Department of Physics, Emory University, Atlanta, GA, 30322, USA; Nonequilibrium Physics of Living Matter RIKEN Hakubi Research Team, RIKEN Center for Biosystems Dynamics Research, 2-2-3 Minatojima-minamimachi, Chuo-ku, Kobe 650-0047, Japan

**Author notes:** **Classification**: Physical Sciences/Applied Physical Sciences; Biological Sciences/Biophysics & Computaitonal Biology.

**Keywords:** epithelial tissue, cell mechanics, cell cycle

## Abstract

Epithelia have distinct cellular architectures, which are established in development, re-established after wounding, and maintained during tissue homeostasis despite cell turnover and mechanical perturbations. In turn, cell shape also controls tissue function as a regulator of cell differentiation, proliferation, and motility. Here we investigate cell shape changes in a model epithelial monolayer. After the onset of confluence, cells continue to proliferate and change shape over time, eventually leading to a final architecture characterized by arrested motion and more regular cell shapes. Such monolayer remodeling is robust, with qualitatively similar evolution in cell shape and dynamics observed across disparate perturbations. Here we quantify differences in monolayer remodeling guided by the active vertex model to identify underlying order parameters controlling epithelial architecture. When monolayers are formed atop extracellular matrix with varied stiffness, we find the cell density at which motion arrests varies significantly but the cell shape remains constant, consistent with the onset of tissue rigidity. In contrast, pharmacological perturbations can significantly alter the cell shape at which tissue dynamics is arrested, consistent with varied amounts of active stress within the tissue. Across all experimental conditions the final cell shape is well correlated to the cell proliferation rate, and cell cycle inhibition immediately arrests cell motility. Finally, we demonstrate cell-cycle variation in junctional tension as a source of active stress within the monolayer. Thus, the architecture and mechanics of epithelial tissue can arise from an interplay between cell mechanics and stresses arising from cell cycle dynamics.

**SIGNIFICANCE STATEMENT:** The morphology of biological tissue is determined by the shape and density of constituent cells. Here we measure the dynamics of cells in model epithelial tissues to study the evolution of their shape and density over time. Guided by a mathematical model, we find that cell shape is controlled by rigidity and active stresses within the tissue. We then show that cell cycle dynamics are the source of active stress that drives epithelial remodeling.

## INTRODUCTION

Cells in epithelial tissues adopt a variety of distinct morphologies which are defined during development and maintained throughout the lifetime of an organism (1). Cellular shape and geometry can be perturbed by stretching or wounding but individual cells within the tissue return to their original shape through increased cellular motility, junctional turnover, neighbor exchange, and proliferation (2–4). In turn, tissue architecture impacts cell fate and tissue physiology (5, 6). Cell division has been implicated as a potential mechanism to regulate monolayer topology (2, 7–10). However, tissue architecture can also change as a result of motion, neighbor exchanges, and shape changes of individual cells in the absence of cell division (11–14). One promising physical framework for predicting collective cell behavior are vertex models, which represent confluent epithelial monolayers by a mechanical network of cell-cell junctions (8, 15–18). From these approaches, mechanical descriptions of epithelial tissue dynamics are being developed (15, 19), but key questions remain.

For instance, it remains unclear what processes set the length scale over which cells in a tissue can move or migrate. This length scale helps determine developmental outcomes, such as whether convergent extension is effective at generating large-scale changes to the body shape, as well disease outcomes, such as whether cells leave a cancer tumor in invasive streams (20, 21). In traditional materials composed of atoms or molecules, particles can freely diffuse when the material is fluid-like, but their motion is arrested in solids, when surrounding particles inhibit their mobility. In the context of biological tissues, it is tempting to speculate that the arrest of motion in a dense collection of cells occurs as the system becomes jammed, or solid-like, which can arise from changes to either density (22, 23) or cell mechanical properties (24–26). However, there is third possible mechanism for arrest of motion: the particles in a fluid-like material could also stop moving if the source of fluctuations becomes very small, despite remaining in a mechanically unstable configuration. Fluctuations in biological tissues are driven by active cellular processes such as cell migration, cytoskeletal contraction, and cell division (27–29). Therefore, the scale of fluctuations in a tissue may be regulated in response to extracellular or intracellular cues to control the degree of tissue remodeling.

In model epithelial tissues, it has been observed that the cells are initially more dynamic – changing neighbors and moving significant distances– and at later times that motion arrests (5, 30–34). Because these changes occur with minimal genetic or biochemical gradients, such epithelial monolayers are an ideal system to study whether cell arrest is governed by an underlying rigidity transition (e.g. collective solidification) caused by changes in density or cell mechanics, or instead by a decrease in active stress fluctuations in a material that remains fluid-like. Particle-based models for tissues predict decreased cell motion arising from reduced interstitial space at increased cell density (12, 22, 23, 35). In contrast, vertex models predict that the cell density is not a direct control parameter for cell dynamics (24). Instead, vertex models predict that observed steady-state cell shape, tuned by varying passive cell mechanics and active forces, is the control parameter for cell motility (8, 17, 19, 24, 25). Cell shape, density and cell adhesion have all been implicated in the arrest of cell motion in epithelial monolayers (5, 30, 31, 33). Furthermore, changes to cell density also regulate signaling pathways that could influence single cell mechanics and cell-cell interactions (36), and cell divisions also introduce active stress fluctuations which in turn affect cell shape (24, 28, 37–39). Thus, it remains unclear how cell density, cell mechanics, and active stress fluctuations contribute to the regulation of epithelial monolayer remodeling dynamics.

Here we use epithelial monolayer remodeling as a model system to investigate the biophysical regulation of epithelial architecture and dynamics. After forming a confluent monolayer, cells continue to divide and change shape over time, until reaching a final state characterized by low motility and more regular cell shapes. Such monolayer remodeling is robustly observed, with qualitatively similar evolution in cell shape and dynamics over a large range of experimental conditions. To tease apart the effects of cell density changes from other mechanical perturbations, we study monolayers formed atop extracellular matrix with varied stiffness. This variation in substrate stiffness causes the cell densities to change significantly, but we find relatively little correlation between density with cell motion. In contrast, we find a striking data collapse when cell velocities are plotted as a function of observed cell shape, as predicted by active vertex models. To understand whether observed cell shapes are primarily regulated by active fluctuations or by changes to single-cell mechanical properties, we perturb the monolayer with pharmacological interventions that interfere with cell proliferation and the cytoskeleton. We find that inhibition of the cell cycle immediately arrests cell motion and shape change, suggesting that cell-cycle-dependent active stress contributes significantly to monolayer dynamics and remodeling. Moreover, across all experimental conditions we find that the average cell shape in the homeostatic final state is well correlated with the cell division rate, suggesting that suppressing cell-cycle-based fluctuations leads to an arrest of cell motion that is independent of the underlying cell mechanics. Finally, we show that cell-cycle-dependent changes in junctional tension are an important source of active stress in the tissue, and use simulations to demonstrate the different tissue architectures that can be realized by modeling cell-cycle-dependent changes in edge tensions. Together our results demonstrate that cell geometry and cell cycle dynamics control cell shape remodeling in epithelial monolayers.

## RESULTS

### Remodeling of confluent MDCK monolayers to achieve homeostatic architecture

To measure the shape and speed of individual cells in a simple epithelial monolayer we created an MDCK cell line that stably expresses green fluorescent protein localized to the plasma membrane via the transmembrane protein stargazin. We seeded these cells at high density on collagen I gels and imaged multiple fields of view using time-lapse fluorescence microscopy. Initially, the monolayer was not continuous, and there were numerous cell-free voids (Fig. 1a). Over time, the cells closed gaps to form a continuous monolayer spanning ∼15 mm; we designate this as t=0 (Fig. 1a). Over the following 12 hours, the cells within the monolayer change shape until a steady-state geometry and density is achieved (Fig. 1a, t=720 min, SI Appendix, Movie S1). Thus, we use this as a model system to study epithelial tissue homeostasis by which cell shape and density is recovered after injury through wound healing and monolayer remodeling (Fig. 1b). While previous work has focused on mechanisms of collective migration in wound healing (40–42), here we focus on the process by which the cells within a confluent monolayer change shape over time.

**Figure 1:**
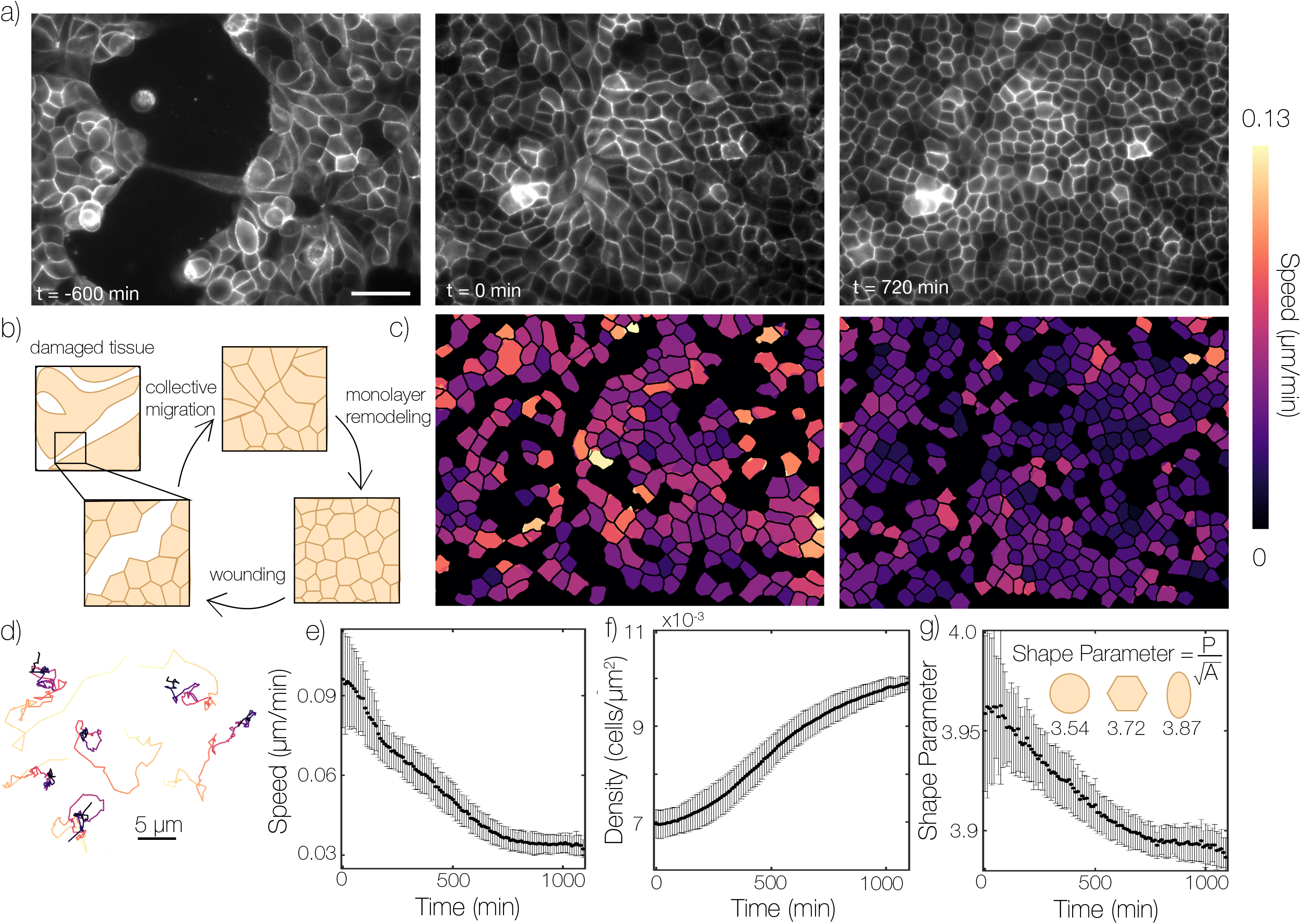
Cell shape remodeling leads to homeostatic monolayer architecture. (a) Cells were plated on collagen gel substrates at t= −1000 minutes. By t = −600 minutes cell have aggregated into large colonies which collectively migrate to fill open space. At t=0 minutes cells have formed a confluent monolayer. Over the next 720 minutes cells become denser and cell morphology becomes increasingly regular. Scale bar is 50 microns. (b) Schematic of the process observed in a). A tissue may be wounded resulting in collective migration followed by monolayer remodeling to the normal epithelial architecture. (c) Heat map of single cell speeds plotted over the segmented cell outlines from images shown in a). Areas in black contain cells which were ignored due to potential segmentation errors. (d) Randomly selected cell trajectories from t=0 to t=1100 minutes, color indicates time. (d-f) Sample averaged values of cell shape, speed and density at each time point across 60 fields of view in the sample depicted in a). Error bars represent standard deviation between fields of view.

We use image segmentation to extract cell shape, size, and positions over time (Methods, SI Appendix, Movie S1). To quantify cell shape, we use cell vertex locations to reconstruct a polygon with a well-defined perimeter *p* and area *A* to calculate the shape parameter (area-normalized perimeter) *q* = *p*/*A*^1/2^; this quantity is bounded from below by objects with a circular shape (*q*_*circle*_ ≈ 3.54), and most cells are observed to have a shape parameter greater than that of a regular hexagon (*q*_*hexagon*_ ≈ 3.72) (SI Appendix, Fig. S1) (24, 25).

We observe cell shape remodeling, along with changes in density and speed, takes place for approximately 12 hours until the system arrests. These dynamics are not sensitive to the details of how time or spatial averaging is performed (SI Appendix, Fig. S2, S3). We also confirmed they are independent of initial seeding density (SI Appendix, Fig. S4), consistent with previous data (31). Similarly to prior observations (30, 34), cell speed decreases (Fig. 1c,d,e, SI Appendix, Fig. S2b) as the density increases from cell proliferation (Fig. 1f). However, we also observe changes in cell shape over time (Fig. 1g). This reduction in shape parameter is correlated with reduced motility in models and experiments where there is no change in density (5, 19, 32). Therefore, from these data alone, the order parameters controlling the steady state cell shape and density in epithelial tissues are impossible to discern.

### Cell shape and Speed are correlated in active vertex models

To further understand the process of monolayer remodeling, we explore predictions of a thermal Voronoi model (43, 44). This model is a variation of standard vertex models which incorporates a simple Brownian noise on each cell to account for active mechanical stress applied by the cells. This model has two key parameters (Fig. 2a): a temperature *T* that represents via uncorrelated noise the magnitude of active stress fluctuations acting on each cell, and a parameter *p*_0_ that represents the target perimeter of each cell and encodes the mechanical properties of a cell, including cell-cell adhesion and tension in the cortically-enriched cytoskeleton. In steady state, these two parameters give rise to a predicted observed shape parameter and cell mobility (19, 44). Importantly, in isotropic tissues a shape parameter of approximately 3.8 reflects the onset of rigidity in the vertex model, whereby higher shape parameters reflect a more fluid-like tissue (19, 24, 25), although the exact location of the transition point depends on the degree of cell packing disorder (45, 46). We expect that the cells tune their active fluctuations and mechanical properties during monolayer remodeling, resulting in potentially time-dependent parameters *T*(*t*) and *p*_0_(*t*) and different “trajectories” through model parameter space (Fig. 2b, SI Appendix, Fig. S5a). Along these parameter trajectories we measure the resulting steady-state shape and speed of cells in simulations (Fig. 2c, SI Appendix, Fig. S5b), which then can be compared to the experimental measurements.

**Figure 2:**
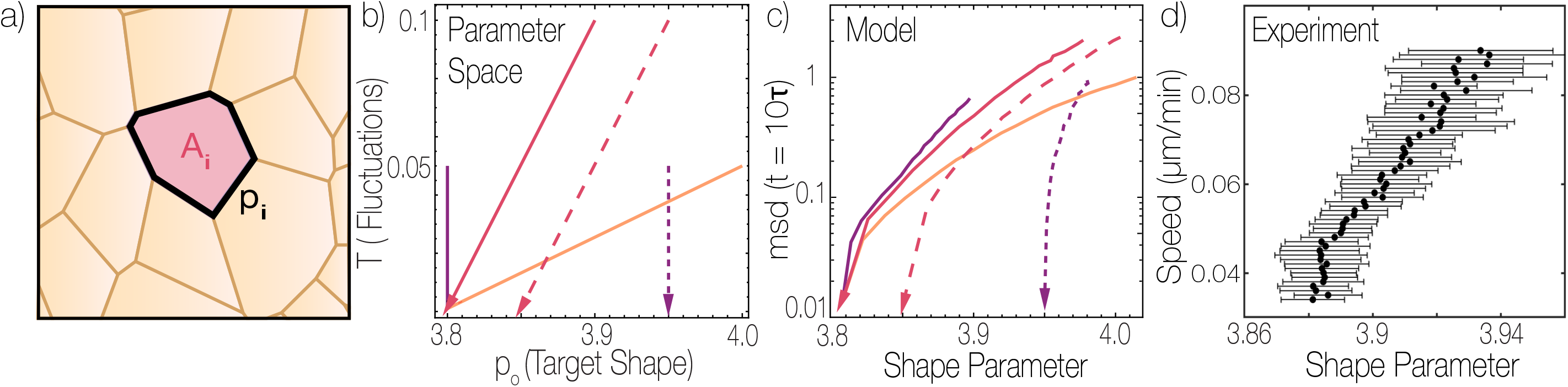
Active Vertex models predict a relationship between cell shape and speed during monolayer remodeling. (a) Schematic of the thermal Voronoi model, where each cell has a target geometry specified by a preferred perimeter *p*_0_ and area *A*_0_. Each cell is subject to Brownian noise with amplitude set by the temperature *T*. (b) Parameters of the thermal Voronoi model were varied along several representative curves. Along these curves a simulated monolayer was equilibrated at each point, after which measurements were made on the monolayer. Solid curves approach T=0 at a value of p_0_ where the tissue is weakly rigid, while dashed curves approach T=0 at a value of p_0_ where the tissue is floppy. Colors represent trajectories with different slopes. (c) Observed values of speed and cell shape. Line styles correspond to the equivalent parameter space trajectories shown in panel b). MSD is given in units of 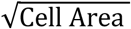 over a time window of 10 natural time units. Data are ensemble averages of 30 simulations each with N=1000 cells. (d) Experimental relationship between cell shape and speed measured for WT dataset on 2mg/ml collagen matrix.

We find that substantially different trajectories through model parameter space can generate similar curves in a plot of typical cell displacements versus observed cell shape, as shown by all of the solid lines in Fig. 2b, c. This is a restatement of the result that, in vertex models, observed cell shape and cell motility are highly correlated (5, 19). In our experiments, we observe a similar relationship between shape and speed resembling these model trajectories (Fig. 2d), suggesting that vertex models may be able to predict features of remodeling in proliferating epithelial layers. Fig. 2b, c also highlights that there is an important exception: the observed shape-motility correlation and convergence breaks down when the temperature (i.e. active stress fluctuation) drops to zero while the cell mechanical stiffness encoded by *p*_0_ is still in the floppy regime, with *p*_0_ ≿ 3.8 (See dashed lines in Fig. 2b). In this case, the “zero temperature” system stops moving, even though the underlying mechanics of the layer is floppy and weak. Since in the real monolayers active stress is generated by cellular processes, we explore these ideas further by considering how monolayer remodeling is impacted by perturbations to extracellular or intracellular pathways.

### Monolayer remodeling is regulated by matrix stiffness and signaling pathways

The stiffness of the extracellular matrix (ECM) can alter cell migration rates through effects on focal adhesion dynamics and cell spreading (47). To explore how epithelial tissues are impacted by the physical properties of the ECM, we varied the underlying collagen gel stiffness and density by either increasing the concentration of collagen, or crosslinking gels with glutaraldehyde (48). This produced gels with Young’s moduli ranging from approximately 200-2000 Pa (49). On all gels, qualitatively similar dynamics in cell shape, motion, and density are observed during monolayer remodeling (SI Appendix, Fig. S6), but there are quantitative differences. For instance, at the onset of confluence when the cell speeds are large (0.07 μm/min), the cell density is ∼50% lower on the stiffer ECM conditions than the soft ones (Fig. 3a, b, SI Appendix, Movie S2). These differences remain as the density increases and cell speed decreases during monolayer remodeling (Fig. 3a, b). Thus, across different matrix stiffness, the number density at which cell speed reaches its minimal value varies substantially (Fig. 3b). This is a strong indication that number density is not directly controlling the arrest of cell motion.

**Figure 3:**
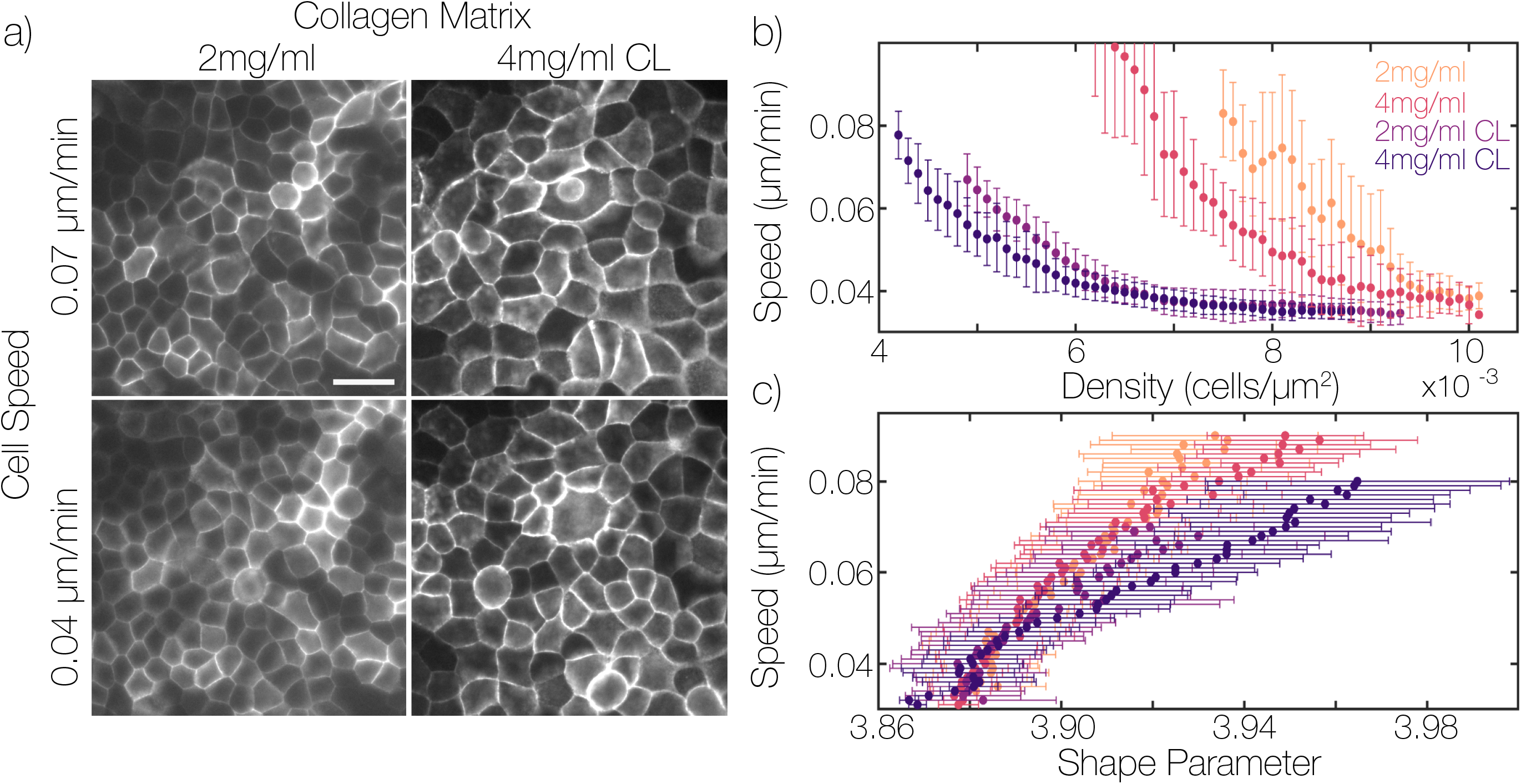
Monolayer remodeling is independent of cell density across perturbations to substrate stiffness. (a) Images of monolayers remodeling on substrates with different stiffness near the beginning and end of the experiment. CL is glutaraldehyde crosslinked collagen gel. Scale bar is 25 microns. (b) Correlation between cell speed and cell density for monolayers on substrates of different stiffness. (c) Correlation between cell speed and shape parameter for monolayers on different substrates. Quantities are averaged over a field of view containing at least several hundred cells for each time point, then field of view measurements are binned together by speed in 0.001 increments. Error bars represent standard deviation of each bin.

In contrast, the cell shapes robustly and reproducibly change during monolayer remodeling, with the arrest of cell motion occurring at a consistent shape parameter of 3.88 (Fig. 3c). Because our collagen gels only span a small range of stiffness, we performed similar experiments on stiffer polyacrylamide gels (16 kPa) and glass and observed similar results, suggesting that this behavior is independent of substrate stiffness across a large range (SI Appendix, Fig. S7). Therefore, cell shape appears to be a robust parameter to predict dynamics and structure of monolayers formed atop different ECM stiffness, in agreement with vertex model simulations.

As demonstrated in the simulation results in Fig. 2b,c, these shape-velocity curves do not by themselves shed light on whether the arrested motion arises predominantly from changes in tissue mechanics or the magnitude of active stresses. An important exception is in regimes where the active fluctuations are driven towards zero while the underlying cell mechanics remains floppy – this scenario results in distinctly different paths through shape-velocity space (dashed lines in Fig. 2c).

To access different regimes of tissue mechanics and active stress generation, we performed a screen of pharmacological perturbations to cell signaling by treatments that altered focal adhesion, cell cycle, and Rho GTPase signaling (SI Appendix, Movie S3, S4). Across all conditions, we observe qualitatively similar monolayer remodeling dynamics that result in arrested cell motion with a characteristic cell shape and density (SI Appendix, Fig. S6, S8). However, there are substantial quantitative changes that contrast with those found in Fig. 3. To illustrate, we consider the impact of fibroblast secreted growth factors by forming monolayers in fibroblast conditioned medium (FCM). We observe that the cell shapes in FCM-treated monolayers are more elongated, with a higher shape parameter, throughout the experiment (Fig. 4a, SI Appendix, Movie S4). At the onset of cell motion arrest, the shape parameter is >3.94 a value much higher than the observed shape in control cells even many hours before arrest (Fig. 4b). This results in significantly higher values of shape parameters throughout monolayer remodeling (Fig. 4b). Across these perturbations, we consistently observe a correlation between the changes in shape parameter and cell speed but observe significant variations in the final shape parameters that occurs at cell motion arrest (SI Appendix, Fig. S8, Fig. S9).

**Figure 4:**
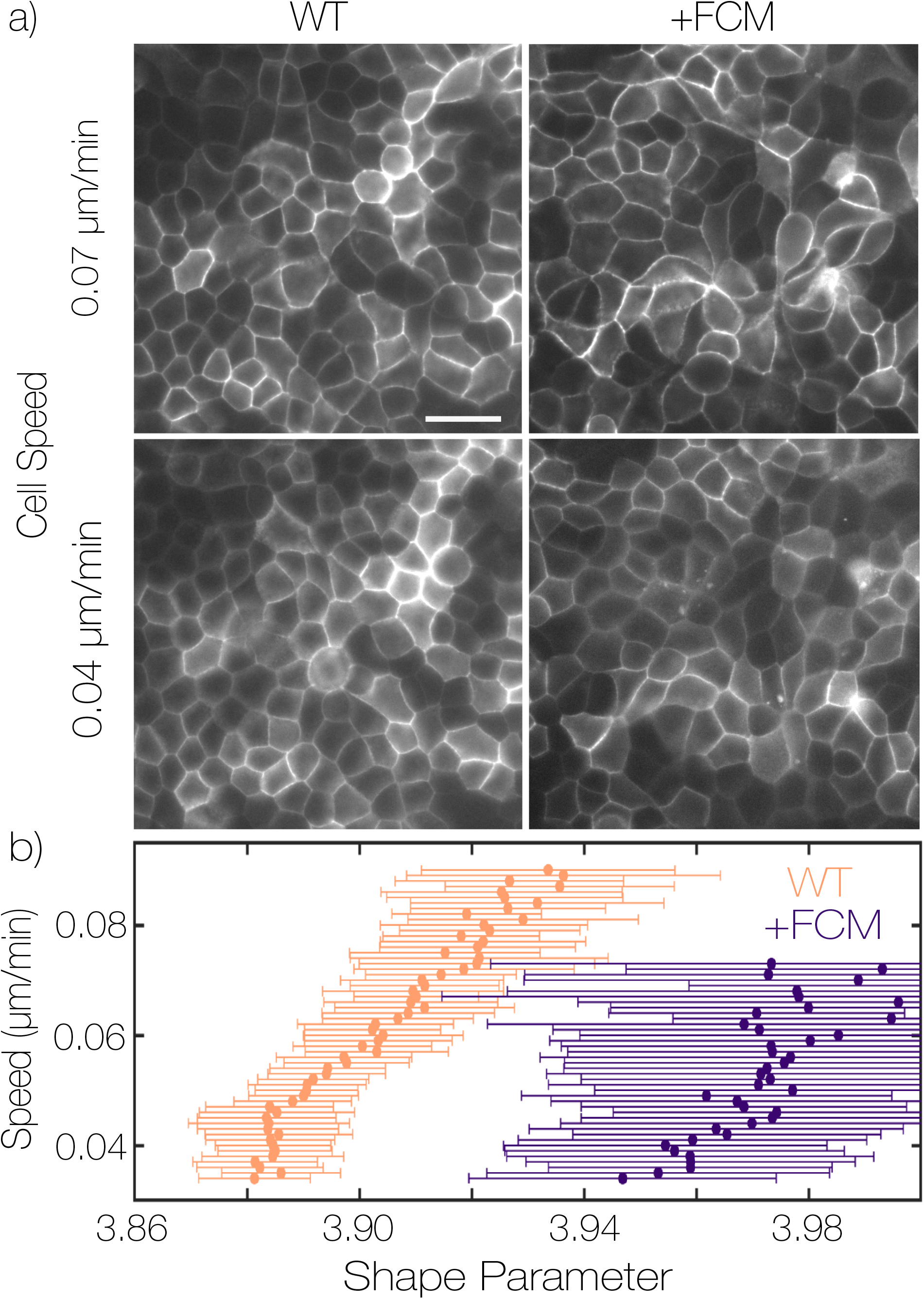
Intracellular signaling alters cell shape during monolayer remodeling. (a) Images of monolayers at the beginning and end of a remodeling experiment under WT conditions or treated with 1:1 fibroblast conditioned medium to culture medium. Scale bar is 25 microns. (b) Correlation between speed and density for control and FCM treated monolayers. Quantities are averaged over a field of view containing at least several hundred cells for each time point, then field of view measurements are binned together by speed in 0.001 increments. Error bars represent standard deviation of each bin.

### Cell division rates control cell shape remodeling

The diversity of monolayer architectures observed across all conditions is demonstrated by plotting the final shape parameters and density (Fig. 5a). With these perturbations, the steady state architecture varies two-fold in density, with cell shapes ranging from elongated to compact, but with no clear correlation between density and shape (Fig. 5a). However, many perturbations reduced the rate of cell divisions (SI Appendix, Fig. S6b). Plotting the final cell shape as a function of the cell division rate for all conditions reveals an inverse correlation (Fig. 5b). Monolayer remodeling obtained from another common epithelial model, CACO-2 cells, with a much lower cell division rate, can be overlaid on this data (Fig. 5b, SI Appendix, Fig. S8).

**Figure 5:**
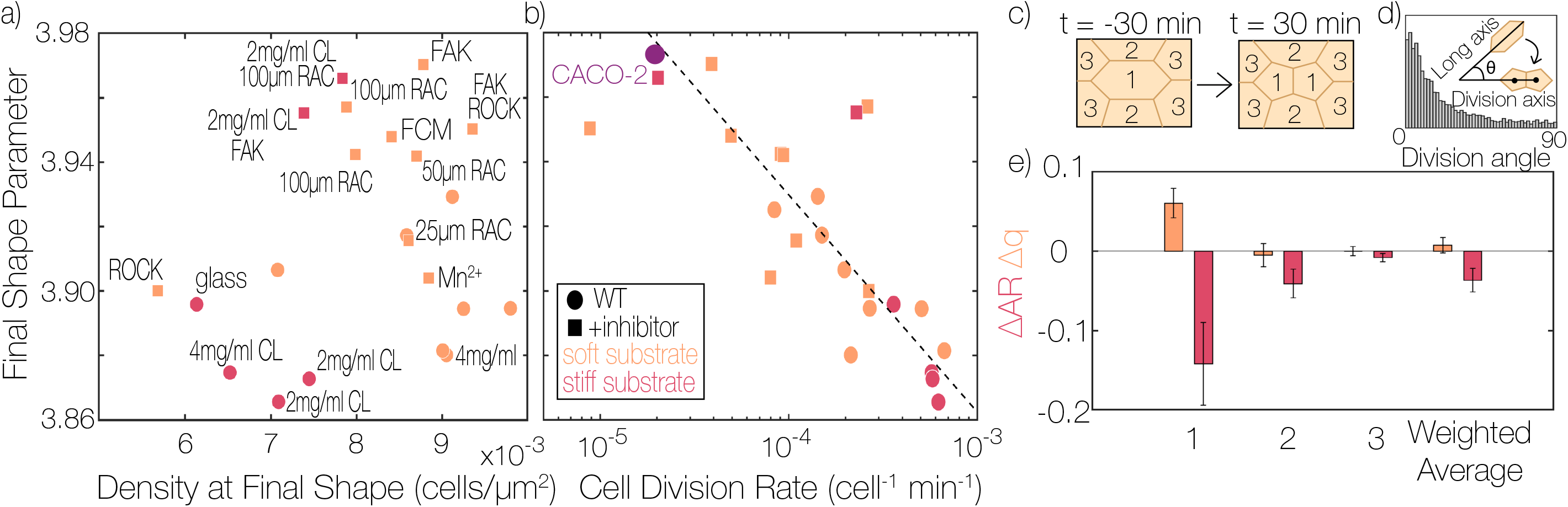
Cell shape remodeling is dependent on cell division rate. (a) Shape parameter vs. density when the monolayer reaches a speed of 0.04µm/min for all inhibitor conditions tested. (b) Final shape parameter vs. the cell division rate from t=200 min to t=600 min for all conditions in a). Logarithmic fit to data is plotted at dashed line. Each data point in a) and b) are the average of >10 time points after reaching the final shape from >30 fields of view from one experiment. (c) Schematic of cell which may experience direct changes in geometry during cell division. 1 is the dividing cell, 2 is a neighbor of both daughter cells, 3 is a neighbor of one daughter cell (d) Histogram of measured angle between the cells interphase long axis and the division plane during WT experiments. (n=3137 cell divisions) (e) Resulting changes in shape and aspect ratio for the dividing cells and neighbors adjacent to one or both cells are plotted. Error bars represent the standard deviation of three WT experiments on 2mg/ml collagen each with at least 1000 cell divisions measured.

To explore how cell division impacts monolayer architecture, we first consider the direct consequences of cell division (9, 10). From purely geometric considerations, one would expect a local reduction in cell shape parameter as a result of topology and aspect-ratio changes after a division (8, 9). Further, oriented cell divisions can directly change cell shape by producing two daughter cells with lower aspect ratio (2, 50). To access the contribution of division to the overall shape change, we analyzed shape change in individual cells in our segmentation data. We measured the shape changes of a dividing cell and its immediate neighbors before and after division, which we classify into three groups based on their contact relationship with the daughter cells (Fig. 5c). Similar to previous findings (2, 7, 50) we observe a strong alignment of cell division along the long axis (Fig. 5c,d). The division results in a reduction of the aspect ratio of the dividing cell but, on average, modestly *increases* the shape parameter (Fig. 5e, Group “1”). Furthermore, there is minimal change in the aspect ratio or shape parameter for neighboring groups of cells (Fig. 5e, Groups “2” and “3”). Considering the weighted average of all groups, the impact of cell division on shape parameter changes is negligible (Fig. 5e, “weighted average”). By tracking shape changes occurring in individual cells throughout interphase, we further verified that the direct effects of cell division are small compared to shape changes occurring through changes in cell junction length in non-dividing cells (SI Appendix, Fig. S10, S11). Thus, local distortions and topological changes during cell division alone are insufficient to explain the contribution of cell division rate to monolayer remodeling.

Other sources of active stresses within monolayers include cell motility (19, 24, 41) and junctional remodeling (51, 52). Compared to other experiments (e.g. wound healing) which show speeds up to 0.7µm/min (40), the extent of cell motion in our experiments is low (<0.1µm/min). One well-documented type of junctional remodeling that drives cell rearrangements is a T1 transition (8), which we observe in the monolayer (SI Appendix, Fig. S12). However, the overall number of cell rearrangements is quite low. Therefore, we surmise that the dominant source of active stresses driving monolayer remodeling is junction length changes in the absence of neighbor exchanges.

### Active stress originates from CDK1-dependent regulation of cell mechanics

Our pharmacological screen included many factors known to impact cell migration, adhesion and force generation, including perturbations to RAC, ROCK, FAK, and Integrin signaling pathways. Surprisingly, these perturbations had little impact on the overall motions within the monolayer (SI Appendix, Fig. S13). However, we did observe a dramatic and immediate decrease in cell speed upon perfusion of an inhibitor of cyclin dependent kinase 1 (CDK1), which blocks the cell cycle. After initiating a monolayer remodeling experiment, CDK1 inhibitor-containing media was perfused in at t=310 min. We observed a striking reduction in cell motion within 20 minutes of inhibiting CDK1 (Fig. 6a-c, SI Appendix, Movie S5). After washing out the inhibitor at t=580 min, there was an immediate recovery of cell motion: remodeling was re-initiated, and the time-evolution of changes in cell shape and speed were similar to those initially observed (Fig.6a-c, SI Appendix, Movie S5). To quantitatively compare these curves, we shifted the post-washout shape parameter data by rescaling the time to (t-400 min); this delay time and rescaling is indicated by dashed lines Fig. 6c&d. Remarkably, with this rescaling the data evolve in almost *quantitative* agreement with each other and, moreover, are comparable to the evolution of unperturbed wild type monolayers (Fig. 6d). This data is consistent with the hypothesis that CDK1 inhibition dramatically reduces the active stress fluctuations (T, in the model). The enhanced motion observed upon inhibitor removal underscores that tuning cell cycle dynamics may be a means to modulate the active stresses.

**Figure 6:**
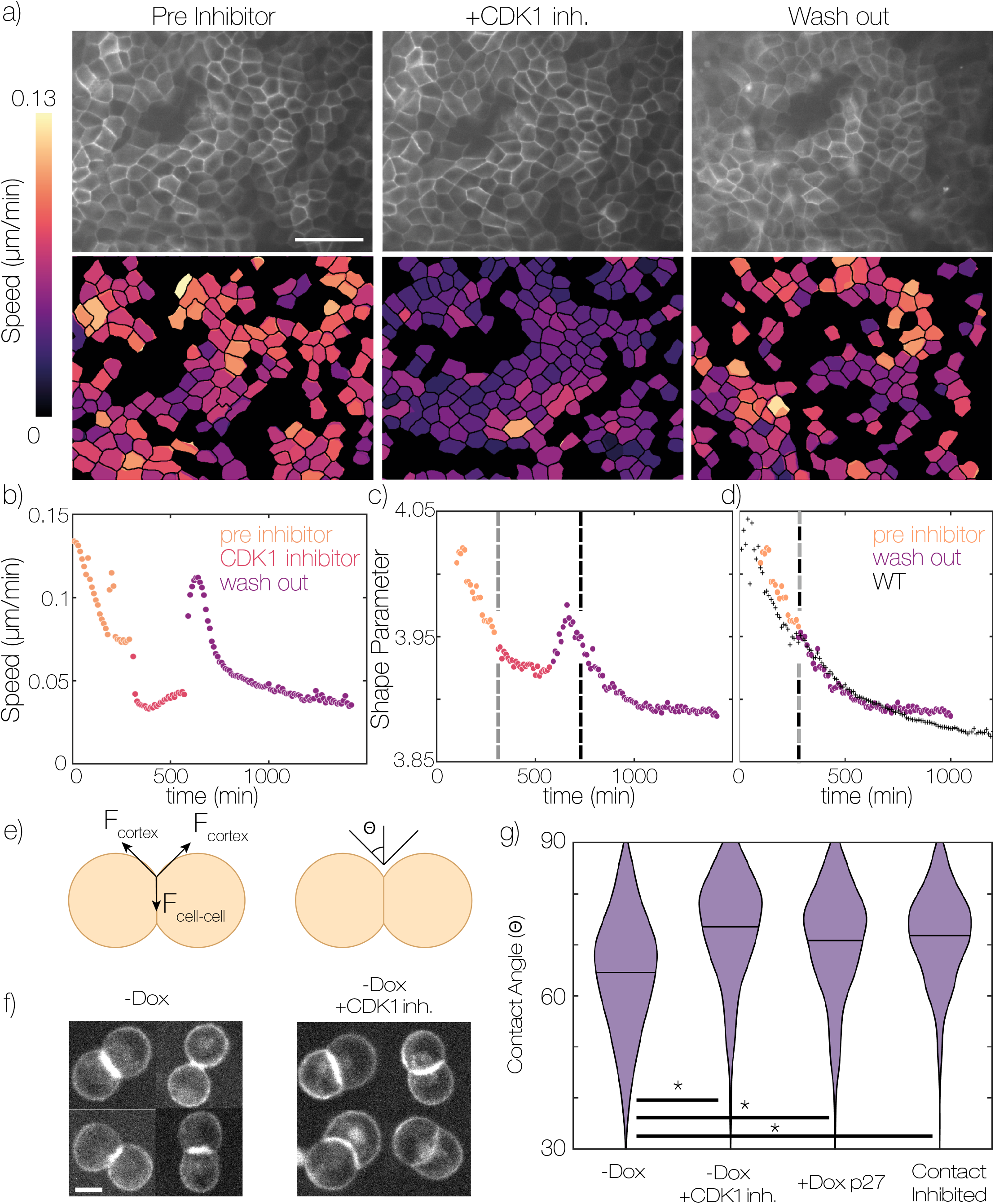
Cell division is a source of active stress required for monolayer remodeling. (a) Representative field of view of monolayer during CDK1 inhibitor wash in experiment. Below each image heat maps of cell speed for successfully segmented cells are displayed. Pre inhibitor is 30 minutes before adding the inhibitor. Inhibitor is 30 minutes after adding the inhibitor. Wash out is 120 minutes after washing out the inhibitor. Scale bar is 50 microns. (b) Average cell speed vs time during CDK1 inhibitor wash in experiment. (c) Average cell shape vs time during CDK1 inhibitor wash in experiments. (d) Time shifted shape vs time curves comparing WT and CDK1 inhibitor wash in data. CDK1 inhibitor wash in data points represent the time average of 12 fields of view in one sample. WT data represents the average of 3 independent experiments each with >20 fields of view. CDK1 inhibitor is 5µM RO-3306. e) Schematic of cell contact angle measurements. Force balance at the contact gives rise to a contact angle Θ. f) representative images of MDCK cell doublets in suspension, in the presence and absence of CDK1 inhibitor 5µM RO-3306. Scale bar is 10 µm. g) Measurement of MDCK tet-p27 cell contact angles under different conditions. -Dox – cells without doxycycline, +CDK1 inh. – 5µM Ro-3306, -Dox p27 – 200ng/ml doxycycline, Contact inhibited – Cells cultured to high density before re-suspension, no doxycycline added. -Dox n=512, -Dox +CDK1 inh. n=480, +Dox p27 n=394, Contact inhibited n=565. Data come from at 3 experimental replicates. * p<0.001

As an alternate means to arrest the cell cycle, we used mitomycin C to abrogate DNA replication, and observed a similar arrest of monolayer movement (SI Appendix, Movie S6). Interestingly, inhibition of cell division by low doses of nocodazole does not reduce cell motion significantly (SI Appendix, Movie S7). Nocodazole allows cells to enter mitosis and initiate mitotic rounding, but prevents further progression of mitosis. This is consistent with our data that cell division, per say, is not the primary source of stress in the monolayer (SI Appendix, Fig. S11). Instead, these stresses arise in the interphase portion of the cell cycle. Together with our characterizations of cell shape changes, we surmise these may come from cell cycle dependent effects on cell mechanics (53, 54).

To test this, we examined the geometry of cell-cell contacts formed by suspended cell-doublets. As demonstrated previously (55), this contact angle Θ can be related to the balance between tension of the cell-cell interface (F_cell-cell_) and cortex (F_cortex_) (Fig. 6e). We first measured the contact angle under both WT and CDK1 inhibitor treatment and noticed a significant increase in the cell contact angles when CDK1 was inhibited (Fig. 6g). To explore whether increased tension at cell-cell contacts is observed by other means of cell cycle arrest, we overexpressed p27^kip^, a protein which binds and inactivates cyclin-dependent kinases, to arrest the cell cycle that is expressed during contact inhibition (56). To isolate CDK-dependent effects, we used a variant of the protein which lacks a c-terminal domain known to interact with RhoA (57, 58). Cell pairs over-expressing p27^kip^ also had increased contact angles, compared to WT conditions. Finally, we measured contact angles of cells obtained from dense contact-inhibited cultures similar to the conditions at the end of monolayer remodeling experiments, and observed an increase in contact angle (Fig 6g). Thus, all of this data demonstrates a cell cycle-dependence of the force balance at the cell-cell interface relative to the cortex. Together with effects of CDK1 inhibition on monolayer remodeling (Fig. 6a-d), these data strongly suggest that cell-cycle dependencies in junctional tension are a primary source of active stress that drives monolayer remodeling.

### Cell cycle arrest leads to low fluctuation arrest of motility in the monolayer

To capture cell-cycle-dependent junctional tension in the vertex model, we built upon previous work that considered the consequences of system-wide fluctuations in interfacial tension (59). Here, we simulate systems where only a subset of edges, which we term “active edges”, generate fluctuating interfacial tension with a characteristic persistence time scale, *τ* (Fig. 7a) (59). At the beginning of these simulations all edges are active, and the fraction of active edges, φ, is reduced from 1 to 0 over the course of a simulation. This emulates the effect of varying the fraction of cell-cycle-arrested cells in the monolayer (Fig. 7a). Concretely, we randomly select an active edge every *τ*^*R*^ natural time units and permanently eliminate the fluctuation of the edge. We observe that as φ decreases the average cell speed diminishes (Fig. 7b). Notably, this parametric plot of speed vs. active edges is nearly independent of the tissue-stiffness parameter *p*_0_(Fig. 7b). To compare this to experiments, we used the pip degron Fluorescent Ubiquitin Cell Cycle Indicator (pip-FUCCI) system to directly monitor cell cycle progression during epithelial remodeling (60). Similar to previous results we see that the fraction of cells in later stages of the cell cycle (S, G2) decreases with time as the cells experience contact inhibition of proliferation (Fig. 7c) (56, 61). When the cell speed is plotted as a function of the fraction of cells which are early in the cell cycle or exited from the cell cycle (G1/G0), we see that an increasing fraction of such cells correlates with a decrease in overall cell motility (Fig. 7d), similar to the simulation results.

**Figure 7:**
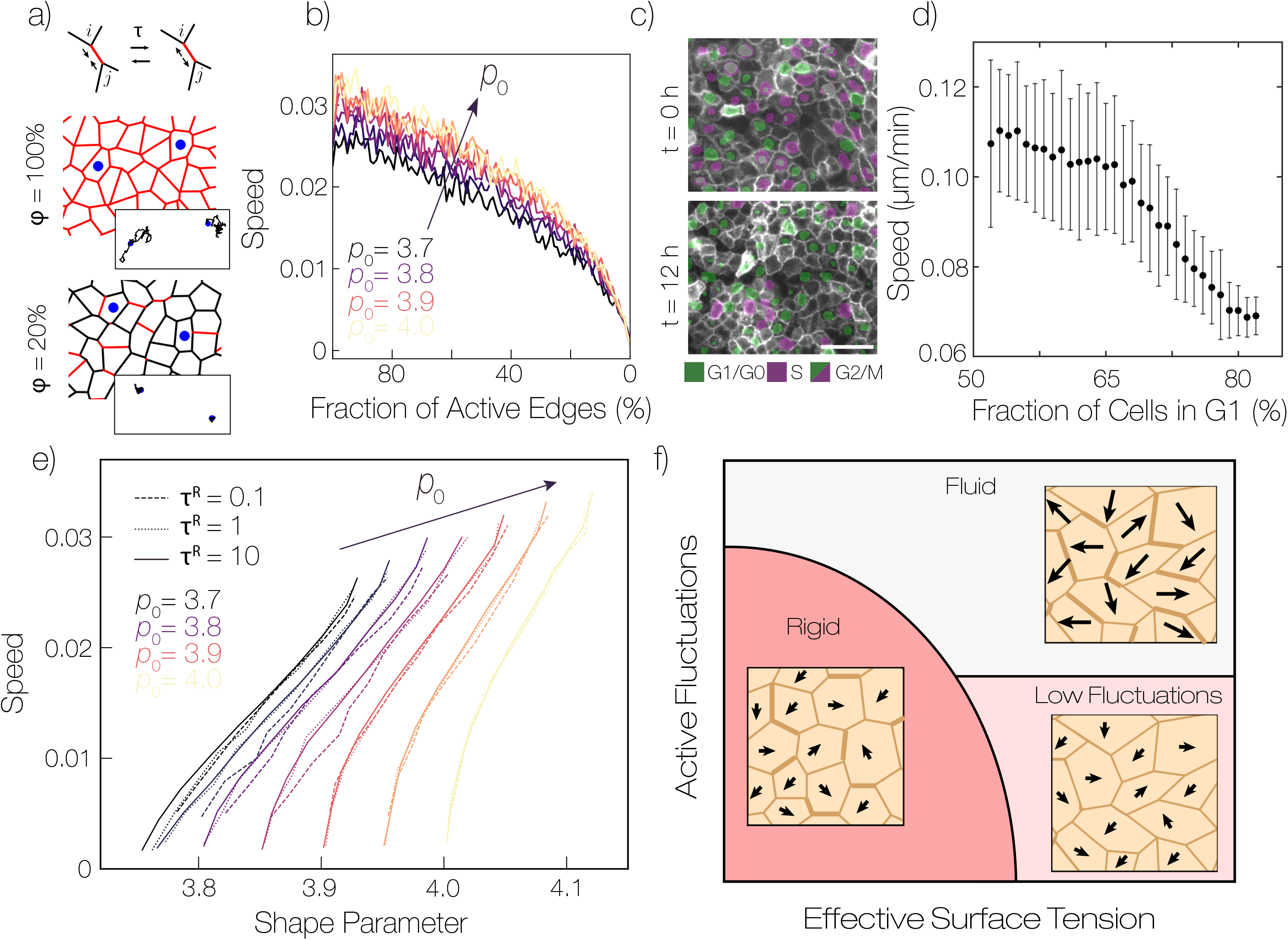
Active Edges as a Source of Stress in Epithelial Tissue. a) Schematic of vertex model with junction tension fluctuations. A fluctuating additional tension is applied between junctions i and j. The fluctuations can be either contractile or extensile, and have a characteristic persistence time *τ* (See the SI for values of all simulation parameters used). Snapshots of cellular configuration at phi=100% and 20% are also shown, respectively (*τ*^*R*^=1). b) Plot of cell speed vs the fraction of active edges in simulated vertex models with fluctuating edge tension (*τ*^*R*^=1). Different curves represent different values of the target shape parameter *p*_0_. c) Images of MDCK monolayers with cell cycle information extracted from pip-FUCCI biosensor. Images from the individual biosensor channels are segmented and overlaid in pseudocolors green (PIP-Venus) and purple (Geminin-mCherry). Presence of only green indicates G1/G0 phase, only purple indicates S phase, and both makers indicates G2/M phase. d) plot of cell speed against fraction of cells in G1/G0 phase of the cell cycle. Error bars represent standard deviation between 30 fields of view. e) Plot of cell speed vs shape parameter from simulations. Curves represent different values of *τ*^*R*^ (dashed, dotted lines) and the target shape parameter *p*_0_ (color scale). f) Schematic phase diagram of epithelial arrest, where edge thickness represents the magnitude of tension on an edge and arrows represent cell displacements.. In addition to fluid and jammed phases which occur at different cell shapes, monolayers can arrest at high average cell shape parameter, indicating a low fluctuation regime. In this low fluctuation regime, the magnitudes of active fluctuations are too small to produce large cell displacements and frequent neighbor rearrangements.

To place this simulation data within a broader framework for cell arrest, we plot the speed versus cell shape for *p*_0_=3.7-4.0. As expected from the homogeneously fluctuation Voronoi model results (Fig. 2), as the fraction of active edges decreases, the speed decreases and the shape parameter approaches *p*_0_ (Fig. S14). Along these curves cell motion arrests at a shape parameter approximately equal to the target shape index *p*_0_, with very little dependence on *τ*^*R*^. Moreover, the qualitative shapes of these curves over a range of *p*_0_ show little sensitivity to the value of *p*_0_ and resemble the experimental data (Fig. 7e). We conclude that at later times in experiments, the cell cycle arrests due to contact inhibition. This leads to a reduction of active stress and the monolayer motility arrests in a “low fluctuation” regime (Fig. 7f). This transition to a low fluctuation regime contrasts with the fluid-solid jamming transition (5, 19, 30). In contrast to the presently proposed scenario, in a jamming transition cell motion would arrest even as large fluctuations in active stresses persist (Fig. 7f). These different scenarios present two distinct paths for controlling cell shape and movement in epithelial tissue.

## DISCUSSION

The mechanisms that regulate epithelial architecture are central to understanding tissue morphogenesis in development, maintenance, and disease. While cell proliferation results in direct changes in topology (9, 10), this does not account for the shape remodeling we observe. In recent years, the development of mechanical models of tissue as active soft materials has provided predictive power to relate local cell mobility and shape. The vertex model predicts that cell motion arrests as cell shapes approach a value that is a sharp rigidity transition, independent of cell density (19, 25). Our data of epithelial remodeling atop matrices with varied stiffness are largely consistent with this model. As the remodeling proceeds, there is 2-fold variation in the monolayer density, but all data collapse on to a “universal” curve of the cell speed as a function of shape parameter. Moreover, the shape parameter at the onset of arrest is 3.88, which is within the range of transition points predicted by vertex models (46). Several previous studied have illustrated the potential of a jamming framework for understanding motion arrest in epithelia (5, 12, 31–33). However, to the best of our knowledge our data is the first to systematically challenge model epithelia with perturbations to demonstrate the robustness of the speed-shape parameter correlation. Importantly, our data strongly supports the utility of cell shape, rather than density, as an order parameter to assert the arrest of cell movement and local epithelial tissue mechanics.

Our data from perturbed systems, however, does not show arrested cell motion at a shape at the predicted jamming transition. Across myriad of pharmacological stimulations and perturbations to signaling pathways, cell motion is arrested even at larger shape parameters and over a wide range of densities. After observing that CDK inhibition immediately arrests the monolayer, we explored the possibility of reducing fluctuations in the monolayer as cells arrest the cell cycle via contact inhibition. In these simulations the monolayer can be arrested at various cell shapes as the fluctuations are reduced. In these conditions arrest occurs at cell shape approximately equal to the model target shape parameter *p*_0_; observing changes in the shape-speed curves corresponds to changing the underlying mechanical properties of the cells. By exploring predictions of the vertex model simulations, a likely possibility is that the effects of these perturbations on active stress are also coupled to changes in the preferred cell shape. However, it may be that additional active-stress-dependent effects on junction remodeling are not captured by our vertex model.

In the fluid regime, the standard vertex model rapidly equilibrates at low temperatures and, at T=0, the final observed and target shape are the same. In such models, perturbations that affect the final shape of cells result in tissues with vastly different mechanical responses. Specifically, these models predict that tissues that arrest due to jamming with target cell shape parameters close to or below 3.8 would be stiffer and less sensitive to small mechanical perturbations than those that arrest due to a decrease in active stress while remaining floppy, with target cell shape parameters significantly above 3.8. Thus, while previous work attributed the arrest of epithelial monolayers to a rigidity transition dependent on density (30, 34) or mechanics (5, 25, 32) our work suggests that arrest can also occur because of a reduction in active stress. Mechanical measurements of epithelia will be required to probe the energy landscape in detail and understand the differences between these scenarios.

In addition, standard vertex models may be too simple to capture the mechanical response in the limit of small active stresses. Recent experiments suggest the existence of dynamic energy barriers, which may prevent equilibration over experimental time scales, arising from junctional stability (4, 62) and/or remodeling (52). In this scenario small but finite fluctuations would not be sufficient to cross those dynamic energy barriers, and so the tissue “freezes” into a metastable state. Thus, a reduction of active stresses could quench the tissue into an arrested state of lower, but non-zero, rigidity. Further work is needed to explore how energetic barriers in remodeling can be exploited for control over tissue mechanics.

The epithelial remodeling we observe is primarily driven by individual junctional length changes, with little contribution from neighbor exchanges or cell division. The limited number of neighbor exchanges observed distinguishes this remodeling from the highly fluid-like behavior observed in other scenarios (51, 63). We find that cell-cycle regulation is the primary source of active stress driving monolayer remodeling. The wide variation in observed shapes across perturbations can be understood by considering their impact on cell proliferation rate. Perturbations that decrease proliferation rate result in motion arrest at higher shape parameters. Moreover, CDK1 inhibition immediately abrogates movement and further epithelial remodeling. While previous data has implicated the importance of mitotic rounding as a source of stress (37, 64, 65) we do not observe large local distortions (SI Appendix, Fig. S11). Therefore, we have demonstrated a role for an active stress generated during interphase or potentially through non cell autonomous behaviors. Cell cycle dependent processes which impact cell adhesion (53), junction tension and cortical mechanics (54, 66) all could give rise to the active stress generation during interphase. Disentangling these effects will be an interesting avenue for future research on cell shape remodeling in epithelia.

Cell proliferation rates can vary widely across different tissues, e.g., the intestine can be entirely replaced on the timescale of days to weeks while the skin may turnover on the timescale of months (67). Moreover, epithelial turnover can be upregulated in response to external stimuli and tissue damage (68– 70). Regulation of cell division rates under these different circumstances may be a way for the epithelium to tune fluidity and facilitate repair. The extent to which cell turnover rate is used to regulate epithelial fluidity and architecture in vivo will be an interesting line of future research. Numerous recent studies have identified a role for cell division in driving morphogenetic processes (13, 64, 71, 72). Our data suggests a mechanism for the increased fluidity observed in highly proliferative tissues through an increase in active stress generation. Measurements of cell shape and motility in these different contexts are required to determine if they are similarly driven by cell cycle-dependent active stress and to discover new mechanisms driving epithelial organization. Further understanding of the interplay between cell cycle, tissue mechanics and cell shape remodeling may lead to a more comprehensive understanding of tissue function in development, homeostasis and disease.

## MATERIALS AND METHODS

MDCK cells were cultured under standard conditions in DMEM with 10% FBS at 37C and 5% CO_2_. Cell monolayers were prepared on by seeding approximately 6×10^5^ MDCK cells on a 500mm^2^ collagen gel substrate overnight. Collagen gels were prepared by polymerizing a neutral collagen solution on silane-glutaraldehye modified glass substrates for 1 hour. Cells were imaged by widefield fluorescence microscopy using standard filter sets. We imaged many locations in the monolayer and verified that they show qualitatively similar behavior (SI Appendix, Fig. S15) Image segmentation, cell tracking and cell division detection were done using custom MATLAB code based on previous methods (73, 74). An example of this segmentation can be seen in the supplement (SI Appendix, Movie S8). By tracking fixed monolayers we found that the tracking errors are small compared to cell motions (SI Appendix, Fig. S16). We also found that stage drift is sufficiently small to be corrected for drift by subtracting the mean displacement. Simulations of the thermal Voronoi model were performed as described in previous publications (43, 44). A detailed description of all methods can be found in the supplementary information.

## DATA AVAILABILITY

Plasmids from this study will be made available from the corresponding authors upon reasonable request and through Addgene (https://www.addgene.org). Code will be made available on the GitHub (https://github.com). Image data will be placed on Dryad.

## ACKNOWLEDGEMENTS

MLG acknowledges funding from NIH RO1 GM104032 and ARO MURI W911NF1410403. MLM acknowledges funding from a Simons Foundation collaborative grant on Cracking the glass problem (#454947) and in the Mathematical Modeling of Living Systems (#446222) as well as NSF-PHY-1607416. TY acknowledges KAKENHI Grant No. 19K16096 and Research Grant from HFSP (Ref. Grant No. RGY0081/2019)/ This project was initiated by Scialog Funding awarded to MLG and MLM.

## Supplementary Information

### Supplemental Information Text

#### Methods

##### Reagents

PND1184, Nocodazole, y27632, NSC23766, Mitomycin C, Human Transferrin, (3-Aminopropyl)trimethoxysilane were purchased from Sigma-Aldrich (Saint Louis, MO), Glutaraldehyde purchased from Electron Microscopy Sciences (Hatfield, PA), BD Collagen I, rat tail was purchased from BD Biosciences (San Jose, CA). 1X PBS, 1X DMEM, Fetal Bovine Serum, l-glutamine, Penicillin-Streptomycin, Trypsin EDTA were purchased from Corning Inc. (Tewksbury, MA), TBS, MnCl, NaOH were purchased from Fisher Scientific (Hampton, NH), Ro-3306 was purchased from Cayman Chemical (Ann Arbor, MI)

##### Cell culture

Madin-Darby Canine Kidney (MDCK) cells and Mouse Embryonic Fibroblasts (MEFs) were cultured in high-glucose DMEM supplemented with 10% FBS, 2mM L-glutamine, 100 U/mL penicillin, and 100 μg/mL streptomycin at 37C and 5% CO2. Caco-2 cells were cultured in high-glucose DMEM supplemented with 10% FBS, 2mM L-glutamine, 100 U/mL penicillin, and 100 μg/mL streptomycin and Human Transferin at 37C and 5% CO2. Cells were passaged using 0.25% trypsin EDTA every 2-3 days. Cells were checked for mycoplasma by Hoechst staining.

Stargazin-GFP MDCK cells we produced by transient transfection of WT MDCK cells with a PiggyBac-stargazin-GFP construct followed by selection by puromycin and subcloning. A clone with high expression of the marker and similar morphology to WT MDCK cells was selected for experiments. Stargazin-GFP was a gift from Michael Glotzer. Stargazin-halotag Caco-2 and MDCK cells were produced by lentiviral infection of WT CACO-2 and MDCK cells by a WPT-Stargazin-halotag construct packaged in 293T cells by a second generation lentiviral system with rev8.2 and VSVG. Viral supernatant was collected at 24, 48 and 72 hours then concentrated ∼30x using Amicon Ultra-15 Centrifugal Filter Unit (100kDa). Cells were treated with virus and 8µg/ml polybrene overnight. Positive cells were isolated using a cell sorter. PIP-FUCCI MDCK cells were produced by lentiviral infection with virus packaged the same way. Cells were then selected using 800 µg/ml G418. pLenti-PGK-Neo-PIP-FUCCI was a gift from Jean Cook (Addgene plasmid # 118616 ; http://n2t.net/addgene:118616 ; RRID:Addgene_118616). Tet P27 1-176 cells were produced using the Lenti-X Tet-On 3G (Takara Bio). Human Snaptag-P27 1-176 was subcloned into the Tre3g vector. Cells were infected with lentivirus for both the Tet-on 3g and Tre3g snaptag-p27 1-176 plasmids then selected using 2 µg/ml puromycin and 800 µg/ml G418.

##### AminoSilane Glutaraldehyde modification of glass coverslips

Glass coverslips were modified as previously described to couple collagen gels to the surface of the glass (Zhu et al., 2012). Coverslips were first cleaned by sonication in 70% and 100% ethanol solutions then dried with compressed air. We placed coverslips in a staining rack and submerged the rack in a solution of 2% (3-Aminopropyl)trimethoxysilane (APTMS) 93% propanol and 5% DI water for 10 minutes at room temperature while stirring. Staining racks were removed and washed in DI water 5 times then placed in a 37C incubator for 6 hours to allow the water to dry and aminosilane layer to cure. The staining racks were then submerged in 1% glutaraldehyde in DI water for 30 minutes while stirring. Then the samples were washed 3 times for 10 minutes in distilled water, air dried and stored at room temperature. Activated coverslips were used within 2 months of preparation.

##### Collagen gel preparation

10x PBS, milli-q filtered water, a 5mg/ml collagen stock and 1N NaOH were mixed to generate a polymerization mix with 1xPBS and 2 or 4 mg/ml collagen at neutral pH. To visualize collagen gel thickness we produced a fluorescently labeled collagen stock by mixing collagen with alexa647-NHS ester in 0.02M acetic acid overnight. We added fluorescently labeled collagen at a 20:1 ratio to unlabeled collagen in the polymerization mix. 70uL of the polymerization mix was added on to a 25mm round modified coverslip and quickly spread to coat the surface using a pipette tip. Samples were transferred to a humidified incubator at 37C to polymerize for 1 hour. After polymerization gels were washed 3 times in 1x PBS and it was verified that gels were still intact and adhered to the glass by a tissue culture microscope.

Glutaraldehyde crosslinked gels were prepared as above and crosslinked as previously described (Lang et al., 2015). Directly after polymerization and washing gels were incubated in 1xPBS containing 0.2% glutaraldehyde for 30 minutes. Gels were then washed quickly 3 times in 1xTBS, washed in 1x TBS at 1 hour intervals 5 times and left in 1x TBS overnight to quench excess crosslinking groups on the gel. The gels were then washed in 1x PBS three times. All gels were used within 2 days of polymerization.

##### Monolayer preparation

Monolayers were formed on collagen I gel to study monolayers under more physiologically relevant conditions than typical glass, plastic or hydrogel surfaces (Bissell, 1981; Elsdale and Bard, 1972; Simons and Fuller, 1985; Walpita and Hay, 2002; Yu et al., 2016). Cells were seeded onto collagen gels at high density (∼600,000 cells on 500mm^2^ surface) such that cells coated ∼70% of the gel surface at seeding. The sample was returned to the incubator overnight (8-12 hours) before the experiment. For inhibitor treated conditions the inhibitor was added 10-20 minutes after the cells were seeded on the gel to allow cells to attach before adding fluid volume. Before mounting samples, the monolayer was viewed under a tissue culture microscope and continuously covered a region 10-15 mm across. Samples were washed 3 times with 1x PBS and quickly mounted into a sealed round chamber with 1.5mL of culture media and with equal concentration of inhibitor to at seeding. Before taking each time lapse the collagen gel was verified by fluorescence microscopy to be homogeneous and 150-250 µm thick. The monolayer was confirmed to extend at least several hundred microns in each direction outside selected fields of view. By only observing cells more than a few hundred microns from a free edge we avoid the increased migration speed and correlation from cytoskeletal assemblies specific to wound healing (Ng et al., 2012; Trepat et al., 2009).

##### Fluorescence microscopy

Cells were imaged on an inverted epi-fluorescence microscope (Nikon TI-E, Nikon, Tokyo, Japan) with a 20x plan flour multi-immersion objective. Glycerol was used as an immersion medium to more closely match the index of refraction of the collagen gel. Images were acquired at 10 minute intervals in GFP, 642 and transmitted light channels using standard filter sets (Ex 490/30, Em 525/30, Ex 640/30, DAPI/FITC/TRITC/cy5 cube) (Chroma Technology, Bellows Falls, VT). Samples were mounted on the microscope in a humidified stage top incubator maintained at 37C 5% CO2. Images were acquired on either a Photometrics Coolsnap HQv2 CCD camera (Photometrics, Tucson, AZ) or Andor Zyla 4.2 CMOS camera (Andor Technology, Belfast, UK).

##### Image Segmentation

Images were segmented using custom MATLAB code. The main algorithm performs initial segmentation using the Phase Stretch Transform algorithm developed by the Asghari and Jalali (Asghari and Jalali, 2015). Phase stretch images were thresholded and skeletonized to obtain cell outlines. Broken edges in the skeleton were repaired using a modified implementation of edgelink developed by Peter Kovesi (“Peter’s Functions for Computer Vision,” n.d.). After edge-linking the remaining unclosed portions of the path were removed. The interior of each cell is checked for high intensity features which typically indicate under-segmentation. Any region containing a high intensity region within the boundary are discarded. The above algorithm has 4 parameters describing the PST parameter set and the interior threshold. Parameters were first optimized by hand. We then randomly generated 1000 parameter sets around these values. We generated a rough ground truth segmentation by averaging over several parameter sets which we verified to segment the images well. The 1000 parameter sets were checked against this ground truth to choose a final parameter set. This parameter set was used to segment all images analyzed in this paper. Examples of the segmented outlines overlaid on the fluorescence images are found in the supplement (Movie S8).

##### Cell tracking

Cell tracking was performed using established particle tracking methods (Crocker and Grier, 1996). Cell centers were determined by taking the centroid of each region in the cell outlines generated as described above in Image Segmentation. The particle trajectories were compiled from these position measurements using SimpleTracker, a MATLAB function developed by Jean-Yves Tinevez (“Simple Tracker - File Exchange - MATLAB Central,” n.d.). For each image in a given time series the average displacement was determined and was subtracted from the cell positions to account for stage drift between each frame. We determined a lower bound on the tracking error by tracking image series of fixed cells (Fig. S15)

For each movie the cell position, shape, magnitude of displacement at a 10-minute interval, and cell area were determined for each individual cell over all time points. The average magnitude of displacement was calculated for each field of view at each time point to obtain a cell speed. The inverse of the average area was calculated to determine cell density in each frame. We computed displacements squared at all subsequent images in the time series and averaged across the dataset to produce a mean squared displacement curve (supplemental figure 3d). Cell positions, velocities and shapes obtained from the tracking were used to compute all correlation functions described in the following sections.

##### Measurement of Cell Shape

We benchmarked a variety of algorithms for determining perimeter and area and found many give rise to large systematic errors. We have chosen to report the shape of a polygon reconstructed from identified cell vertex locations which we found to be the most accurate and robust metric (Fig S1, supplemental discussion). Shape parameter was computed for each cell based on the perimeter and area of a polygon constructed from cell vertex locations. The vertex locations parameterize the vertex model and therefore also allow for direct comparison with the model. Reconstructing a polygon from vertices also removes resolution dependent ambiguity in perimeter measurement (Coastline Paradox).

Vertex locations were found by locating branch points in the segmentation mask generated as described in Image Segmentation using the built in Matlab bwmorph function. Cells which do not have a complete set of segmented neighbors also do not have a complete set of vertices and therefore were discarded. We measured average cell shape for the full set of cells and cells remaining after discarding edge cells and found only a small difference in the average. We believe this difference comes from a small segmentation bias for cells whose outline is not correctly constrained by the neighbor. Therefore, the interior cell shapes are likely slightly more accurate. The shape parameter q = perimeter/sqrt(area) was measured by computing perimeter and area for the reconstructed polygon. The shape parameter is related to the another popular shape metric – circularity = 4*pi*area/perimeter^2^ = 4*pi/q^2^. Final cell shape was determined for each experiment by averaging the shape parameter for all fields of view measured to have a speed below 0.04µm/min. This final shape value is within a few percent of the value obtained by fitting the shape vs time curves to an exponential decay.

##### Simulation methods

We perform numerical simulations of monodisperse thermal Voronoi models, as described in (Sussman, 2017). Briefly, we begin by writing down a dimensionless form for the standard vertex model energy, 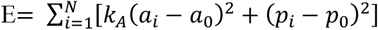. This simple expression assigns an energy to the cells in a confluent monolayer in terms of their preferred geometry. The energy depends on the area *a*_*i*_ and perimeter *p*_*i*_ of each of the *N* cells (indexed by *i*), which are determined by a Voronoi tessellation of the cell positions. The unit of length in the simulations are defined such that the average cell area is unity, and we also set both the preferred area *a*_0_ *=* 1 and the stiffness parameter *k*_*A*_ *=* 1. The preferred value for the cell perimeter, *p*_0_, then constitutes the remaining control parameter which sets the target state of the monolayer.

We then use the cellGPU package to simulate overdamped Brownian dynamics of the model at different temperatures, *T*. The curves in Fig. 2c were created by performing an ensemble average over approximately 30 independent systems of *N =* 1000 cells at each (*p*_0_, *T*) point in parameter space along the lines indicated in (Fig. 2b). Each system was initialized in a high-temperature configuration and then allowed to equilibrate at the target temperature for a large multiple of the system’s characteristic relaxation time, as estimated from the data in Ref. (Sussman et al., 2018); after this equilibration period the observed average shape parameter of the cells and mean-squared displacement at several typical time lags was evaluated.

Numerical simulations of the vertex model with fluctuating junctional tension are performed as previously described (Yamamoto et al., 2020). The dimensionless energy of the vertex model is written as a function of the vertex coordinates 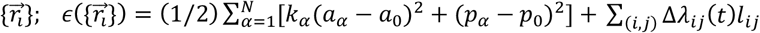. Here, *α* and *N* denote the label of each cell and the total number of cells, *a*_*α*_ and *p*_*α*_ are the area and perimeter of cell *α*. The preferered area and perimeter are denoted as *a*_0_ and *p*_0_, respectively. *k*_*α*_ is the stiffness parameter. In the non-dimensionalized form, we choose the length scale to satisfy the average cell area ⟨*a*_*α*_⟩ = 1. The edge length between the vertices *i* and *j* is denoted as *l*_*ij*_, and the fluctuating line tension is introduced by Δ*λ*_*ij*_(*t*). The dynamics of Δ*λ*_*ij*_(*t*) is described by an Ornstein-Uhlenbeck process satisfying ⟨Δ*λ*_*ij*_(*t*)⟩ *=* 0 and 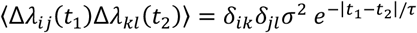. The vertex dynamics is described by the time-evolution equation 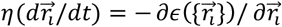. In this paper, we choose the *σ =* 0.2, *τ =* 1, *k*_*α*_ *=* 1, *a*_0_ *=* 1, and *η =* 1, and we run simulations with 340 cells in a square simulation domain with periodic boundary conditions. As we for our Voronoi model simulations, we initialize the cellular configuration under a high-temperature condition (*σ =* 0.35, *τ =* 1), and thermalize with the target parameters. After the thermalization, we begin setting the tension Δ*λ*_*ij*_(*t*) of edges to zero permanently by randomly selecting a target edge every constant time interval *τ*^*R*^.

##### Preparation of Fibroblast conditioned Medium

Fibroblast conditioned medium was prepared as described previously (Montesano et al., 1991). We cultured a 10 cm dish of MEF cells to confluence. Cells were washed and 20 mL of fresh medium was added to the cells. 48 hours later the media was collected and centrifuged to remove any cells. Conditioned media was aliquoted and frozen at −20C then used within 1 month of preparation. The conditioned medium was mixed 50:50 with fresh culture medium and added to MDCK cells 15 minutes after plating. We tested that fibroblast conditioned medium caused cell scattering of small colonies on collagen coated glass substrates as previously described (Stoker et al., 1987).

##### Cell division tracking

We found that our cell tracking consistently does not follow cell trajectories through a cell division because the distance threshold of trajectory linking is significantly larger than a cell radius. We exploit this to identify cell divisions and to measure shape change independent of cell division (Fig. 5d, Fig. S10). We identified cell divisions by identifying pairs of cells which appear in adjacent to each other in a frame after both cells were not present in the previous frame. We further filter out cells which are not of similar size to one another. We confirmed by inspection that this gives us a subset set of cells which have divided in the previous frame with few false positives. We then find the mother cell by looking several frames back for a cell near the centroid of the pair of daughter cells. We identify neighbors adjacent to both or one of the daughter cells and track each of these cells back in the trajectory to compare cell shapes before and after the division. We then average across all cell divisions in the dataset to produce the final values (figure 5e).

##### Calculation of mean squared displacements

Particle trajectories were compiled as described in the Particle Tracking section of the methods. We took each trajectory and decomposed it into non-overlapping sub trajectories starting from the initial time point with length ranging from tau = 10 minutes to the full trajectory length. The average displacement squared for each sub trajectory was computed and averaged across the dataset (Fig. S5d). To compute the time averaged mean squared displacement averaging was done for all sub-trajectories of a single particle and each cell with a full trajectory is plotted (Fig S2a). To plot mean squared displacement against time we averaged for a given value of tau and t across the dataset (Fig. S2b).

##### Three Pixel vector method of measuring perimeter and area

The three pixel vector method was implemented as previously described (Inoue and Kimura, 1987). Briefly, the different ways of linking pixels are divided into 4 classes and 13 subclasses. Each subclass has a defined length giving a 1×13 vector. For each class there is a linking correction depending on the difference of the direction of the current and subsequent vector which is specified by a 4(classes)x16(directions) matrix. To determine the areas enclosed by each vector we sketched the set of 13 available classes of linking and computed the area of edge pixels and pixels outside the polygon for each case. We obtained the values [2,3,2,2,2,2,3/2,1,11/4,3,2,3/2,2]. We then inferred area corrections for each class from their respective perimeter corrections linking as follows : [0,0,0,0,1/2-2,1-1/2,2-2,0,2,2,2,2,1,1,1,1]; [0,0,1/2-2,1/2-2,2-5,0,0,2,2,2,1,1,1,1,0,0]; [0,-1,-1,2-2,0,0,0,0,2,1,1,1,1,0,0,0]; [0,0,0,1/2-2,2-2*1/2,2-2,0,2,2,2,2,1,1,1,1,0]. We compute the perimeter as described by Inoue et al. and using the same rules and values specified above plus the area of the interior pixels calculate the area of the polygon.

##### Field of View analysis

The shape vs speed for different size fields of view were computed by segmenting each field of view into a set of sub fields. To simplify this process first the field of view was truncated into a square to make sub division possible in such a way that the same set of cells are always measured. Then the field was subdivided into 1, 2×2, 3×3… 10×10 regions. We computed the average speed and shape in each sub-region, then binned the results of all regions according to average speed in the region. To characterize noise in the correlation we computed the average deviation of the derivative in a linear region of the curve for shape parameters ranging from 3.93 to 3.97.

#### Measurement of neighbor exchange rate

To measure a neighbor exchange rate, we detected 4 fold vertices and computed the ratio of 4 fold to 3 fold vertices per unit time. Such 4 fold vertices are not stable in the system and thus result in either a successful or attempted neighbor exchange. Our method does not detect the difference between successful neighbor exchanges and attempts which resolve in the original direction. A 4 cell vertex after formation was observed to resolve in either direction within at most 10 frames. Therefore, we iterated through each set of outlines and detected all 4 cell vertices. We discarded detections which happened within 10 pixels and 10 frames of a detected event to avoid double counting due to time delay between formation and resolution. This set of candidate events contained many false positives where the outline appeared to have a four-fold vertex but upon inspection of the raw data we observed a short 3 cell vertex. To remove these false positives, we manually sorted through the 4 cell vertex candidates and selected out real events.

#### Cell Doublet preparation and measurements

Cell doublets were produced by treating cells with trypsin until detachment from cell culture dishes. Around 15,000 cells were transferred to a PDMS well which was pretreated with 1% Pluronic f127 for 1 hour then washed 3 times with 1X PBS. Cells were incubated in the wells at 37C and 5% CO2 for 4 hours in the presence of inhibitors or doxycycline (100ng/ml) and halotag and snaptag ligands (1µM) then imaged at room temperature for less than 30 minutes. For the “Contact inhibited” condition cells were plated at nearly 100% confluence and grown for 3 days prior to detachment while other conditions were cultured under normal cell culture conditions. Cells were imaged by mounting PDMS wells on coverglass and using imaging methods described above. The contact angle was measured between isolated pairs of cells manually using imageJ.

#### FUCCI Measurements

Cells were imaged in GFP and RFP channels similar to above methods. Images of FUCCI markers and cell boundaries were segmented using Phase Stretch Transform in Matlab. Each cell was identified using the cell boundaries and was determined to be GFP or RFP positive by measuring the intensity contained within the segmented images of each nuclear marker. The percent of cells in G1 was determined by taking the ratio of cells identified as only GFP positive to the cells identified as GFP positive, RFP positive and positive for both markers.

## Additional Discussions

### Shape Metric Benchmarking - Common Shape Metrics are resolution Dependent

In the initial analysis we noticed an unexpected density dependence of cell shape which had a different magnitude depending on which metric we used for measuring shape. We therefore decided to benchmark methods of measuring object shapes in digital images. Measuring the shape of polygonal objects projected onto a pixel grid is nontrivial because segmented edges are limited to single pixels while the actual object edge is a subpixel feature. Data is recoded on a 6.45µm/pixel camera at 20x magnification giving 0.3225 µm/pixel final resolution for a cell radius of ∼15µm. Therefore, the edge of a cell contains few enough pixels that choice of shape metric is important. The simplest method to determine a length in an image made of pixels is to count the number of pixels along the perimeter. A more sophisticated method count pixels at 0, 90,180,270 degree angles (even) as a distance 1 and diagonal (odd) pixel connections as sqrt(2). This method is used to determine perimeter in the popular image processing software imageJ. This method overestimates the length where even and odd edges meet. Another method developed by Vossepoel uses a correction factor for pixels which change from odd to even with parameter values fit from simulated images of lines. This is used in the built in perimeter determination in regionprops in Matlab. This correction factor is not always accurate for small polygons. The corner correction can also be made by estimating that these connection have length sqrt(5) although this method does not correctly capture the length of some corner corrections. The Three Pixel Vector method developed by Inoue and Kimura explicitly implements all possible corner corrections (Inoue and Kimura, 1987). Object areas also can be measured in several ways. The most common method is to add up all the pixels in the polygon which the default method used in ImageJ and Matlab. This method overestimates the area of the polygon as it encloses area outside of the corrected perimeter contours described above. Instead the area can be measured by constructing the contours described above and finding the enclosed area.

To measure shape we implemented several of the methods described above. We also implemented an additional method which is specific to our dataset, where the objects can be defined by simple polygons constrained by the cell vertices. To measure this polygon shape we measure vertex locations estimated to the nearest pixel and reconstruct the polygon by connecting the vertices by straight lines. The vertex locations were determined by locating branch points in the cell outlines. One drawback of this method is that segmented cells which do not have a complete set of neighbors lack a complete set of vertices and must be discarded. For every cell with a complete set of neighbors, the vertices belonging to each cell were determined. The perimeter and area of a polygon defined by a set of vertex locations can be computed directly without lines being interpolated on to a grid of pixels. A representative image of the vertex locations on the membrane GFP image are displayed (Fig. S1b).

We compared the relationship between cell shape and density which reveals the presence of resolution dependent artifacts at increased density (Fig S1c). We observe a larger density dependence for more simple methods of perimeter and area determination which are known to produce large resolution dependent errors. The methods from three pixel vector closely match the vertex reconstruction – we infer that these are the two most accurate methods. We confirmed these findings by generating simulation images from the thermal voronoi model with known shape parameter and measuring the cell shapes (Fig S1d). This confirms that TPV and vertex reconstruction typically have less than 1% error. Finally, we segmented images of fixed cells (Fig S9) and measured the cell shape over a trajectory. This shape should be constant because there is no motion in the images only small changes in signal to noise and small stage drifts. We compared the measured values across each trajectory to the mean and observe larger fluctuations in the TPV method (Fig S1e). This indicates that small differences in the segmentation boundaries can lead to relatively large errors in the final value of shape parameter. Therefore, the vertex reconstruction appears to be more robust to small changes in the signal. Although the vertex reconstruction requires discarding cells with an incomplete set of neighbors, for our 335×445 µm field of view and signal to noise the number of measurements (100-400 cells) is still large enough to provide a reliable average.

### Cell Motility decreases with time

We observed that dynamics in the MDCK monolayers evolve with time. To confirm that individual cells show similar motile behavior we plot the time averaged mean squared displacement (TAMSD) (supplemental figure 2a). We observe qualitatively similar behavior for all cells in the monolayer. At short time scales the motion is nearly ballistic, however at later lag times the TAMSD plateaus for each cell. This indicates that at longer time scales most cells are confined by the neighboring cells and as a result do not end up traveling more than a few microns – only a fraction of a cell diameter. We also look at the time evolution of this motion by plotting the mean squared displacement as a function of time (supplemental figure 2b). We observe that for different lag times the mean squared displacement decreases. This shows that at later time points even for short time scales the diffusion is slower.

### Correlation between shape and speed is not dependent on field of view size or lag time

From our vertex model, we expect that shape parameter is useful for describing dynamics in the system. In the model, passive and active forces are defined at the level of single cells and therefore we expect that behaviors are mainly dictated by a cell and its nearest neighbors. The relationship between shape and speed should not be dependent on how the system is measured, meaning that the size of the field of view or lag time is somewhat arbitrary. We first test the relationship between the correlation we observe and the field of view size. We subdivide the field of view into at 335µm square, 2×2 168µm squares, … 10×10 35µm squares (Fig. S3a). These squares contain the same data but if the properties of the system were not uniform, the smaller partitions may show different behaviors. If the forces are defined at the cellular scale, as we expect from the model, this partitioning should not change the relationship between shape and speed. We observe that across all partitions the same final relationship is recovered (Fig. S3b). However, we see that there is less noise in the correlation curve when partitioning into smaller regions. Because the correlation curve is not linear and the field of view is not uniform, as we average over a larger area the average shape does not capture local shape variation which results in regions much faster or slower than the average. We plot the fluctuations in the correlation curve with respect to the average to describe this effect (Fig S3c). As we reduce the size of the region we see the curve becomes more smooth until we reach a noise floor at ∼100µm. We anticipate that if we had lower noise of cell displacements and a full segmentation these fluctuations would continue to decrease to the scale of a few interacting cells.

We also wanted to ensure that this correlation is not dependent on the lag time within a range of timescales. We plot the mean squared displacement as a function of the shape parameter at the beginning of each trajectory (Fig S3d). We see that for each lag time there is still a relationship between shape and speed. We plot the relationship between shape and speed for one field of view at different lag times and see similar behavior (Fig S3e). The relationship shifts down because the motion is diffusive so as lag time is increased by a factor of two displacement increase by a factor of sqrt(2). We also see the same trend for the ensemble averaged correlations (Fig S3f).

### Cytoskeletal rescue experiments do not restore relationship between shape and speed

Across inhibitors tested we examined the relationship of shape and speed. A subset of these conditions is plotted (Fig S8). In general, we see that the relative relationship between shape and speed for a given condition is similar however the absolute values are all shifted in the inhibitor cases. One potential concern in these experiments is that cytoskeletal polarization may be perturbed (Akhtar and Hotchin, 2001; Bryant et al., 2014; O’Brien et al., 2001; Playford et al., 2008; Yu et al., 2003). Changes in cytoskeletal organization may affect how well the system is described by vertex models which assume the monolayer mechanics are dominated by cortical actin at the cell-cell junctions. We wanted to examine if these defects are related to changes in cell polarity observed for many of these inhibitors. We attempted rescue experiments based on published rescues of RAC and FAK knockdown experiments. It has been observed that RAC promotes apical basal polarity through its role in assembling laminin into the basement membrane (O’Brien et al., 2001). We seeded cells on a collagen gel with 1mg/ml matrigel to see if the inclusion of exogenous laminin in the matrix via the matrigel would rescue the relationship between shape and speed (Fig S9b). We observe the same relationship between shape and speed in the presence of matrigel. We also attempted inhibiting both FAK and ROCK at the same time. ROCK inhibition rescues wound healing in FAK knockdown cells and restores apical basal polarity (Bryant et al., 2014; Chen et al., 2002). We observe similar behavior in conditions with both inhibitors (Fig. S9a)

### Neighbor exchanges rates are low

We measured the rate of neighbor exchanges for one wild type and one inhibitor dataset (Fig. S12a). We were not able to reliable measure neighbor exchange rates with automated analysis so we only made this measurement for these two datasets. These neighbor exchange rates range from 1-10 per hour per 1000 cells which seems low compared to observations from some phases of development where neighbor exchanges are important for tissue flow (Blankenship et al., 2006). We observe that neighbor exchange rates are shape dependent and lower in the inhibitor treated conditions at all shape parameters. This suggests that the rate of neighbor exchange also depends on the cell division fraction, or active stress in the system. We relate these neighbor exchange rates to observed velocity and observe a correlation (Fig S12b).

### A metric q_track_ can be related to different modes of shape change

We wanted to confirm that the relationship between final shape and cell divisions was not simply the consequence of oriented cell division. We exploit the fact that q_track_, the difference in cell shape between subsequent time steps (q_track_(t) = <q(t+1)-q(t)>), measures certain forms of cell shape change while ignoring others. For a given time step we directly observe, by tracking, a subset of the shape change which occurs. We denote the two types of cell shape change, division based (Q2) and deformation based (Q1 / q_track_). The observed subset selectively ignores cell divisions, because division results in large enough displacements that the trajectory is broken in our tracking algorithm. q_track_ also ignore a subset of shape changes where the cell is not segmented in the initial or subsequent frame which are of either type (Fig S10a). Importantly, only type 1 shape changes are observed in q_track_. Therefore, as long as there is not a difference between the behavior of cells which are not segmented we can obtain the average value of a Q1 shape change for a give frame. We can also determine the total shape change by measuring shape in each frame over all cells. We show that the ratio of these two, multiplied by a small correction factor (∼1%), gives a relationship for the relative value of non-division based shape change (Fig S10b, S10c). If this value were on average 0 we could explain the correlation in figure S10d by oriented cell division. We observe values much larger than zero for this metric demonstrating there is an additional mechanism driven by cell divisions. Our calculation ignores the addition of a neighbor to cells adjacent to the dividing cell. We confirm that the shape change by q_track_ is several times larger than the shape change which would result from the gain of an additional neighbor (Fig S10e). We observe that this metric is nearly 1 across all average shape parameters for three datasets with very different division rates (Fig. S10e). We show that for one condition q_track_ is similar to the total shape change except at early time points (Fig. S10f). We also observe that the direct shape change before and after cell division is shape dependent in this dataset consistent with the deviations in q_track_ we observe at high shape parameter (Fig. S10g).

### q_track_ Derivation

The following derivation shows how the ratio of q_track_ to q_total_ can be interpreted in Fig S10. We see that the ratio is 1 if shape change through division is small or zero if shape change through division is equal to the total.

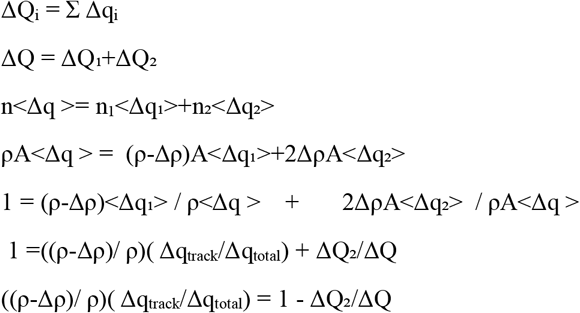

### q_track_ values are consistent with junction length changes as the main source of shape change

We measured shape change along cellular trajectories Δq_track_(t) = <q(t+1)-q(t)> which we show measures deformation based changes and ignores oriented division effects (Fig. S10b; S10c). We then compare this to the total shape change between time points Δq_total_(t) = q(t+1)-q(t) to measure the relative contribution of junction length changes and oriented division (Fig. S10e). We observe a slight discrepancy between these two metrics at early times in the experiment. At these time points cell aspect ratios may be large enough for oriented division to cause a net decrease in cell shape (Fig. S10d). However, we observe at later time points the q_track_ metric which ignores oriented division effects is sufficient to capture nearly all shape change in the monolayer. Across conditions with different division fraction junction length change consistently explains a majority of shape change in the monolayer (Fig. S10e). Therefore, the relationship in figure 5d implies that in monolayers with higher division fraction there is additional cell shape remodeling which occurs as a result of differences in cell mechanical properties or active stress caused by cell division.

---

**Fig. S1:**
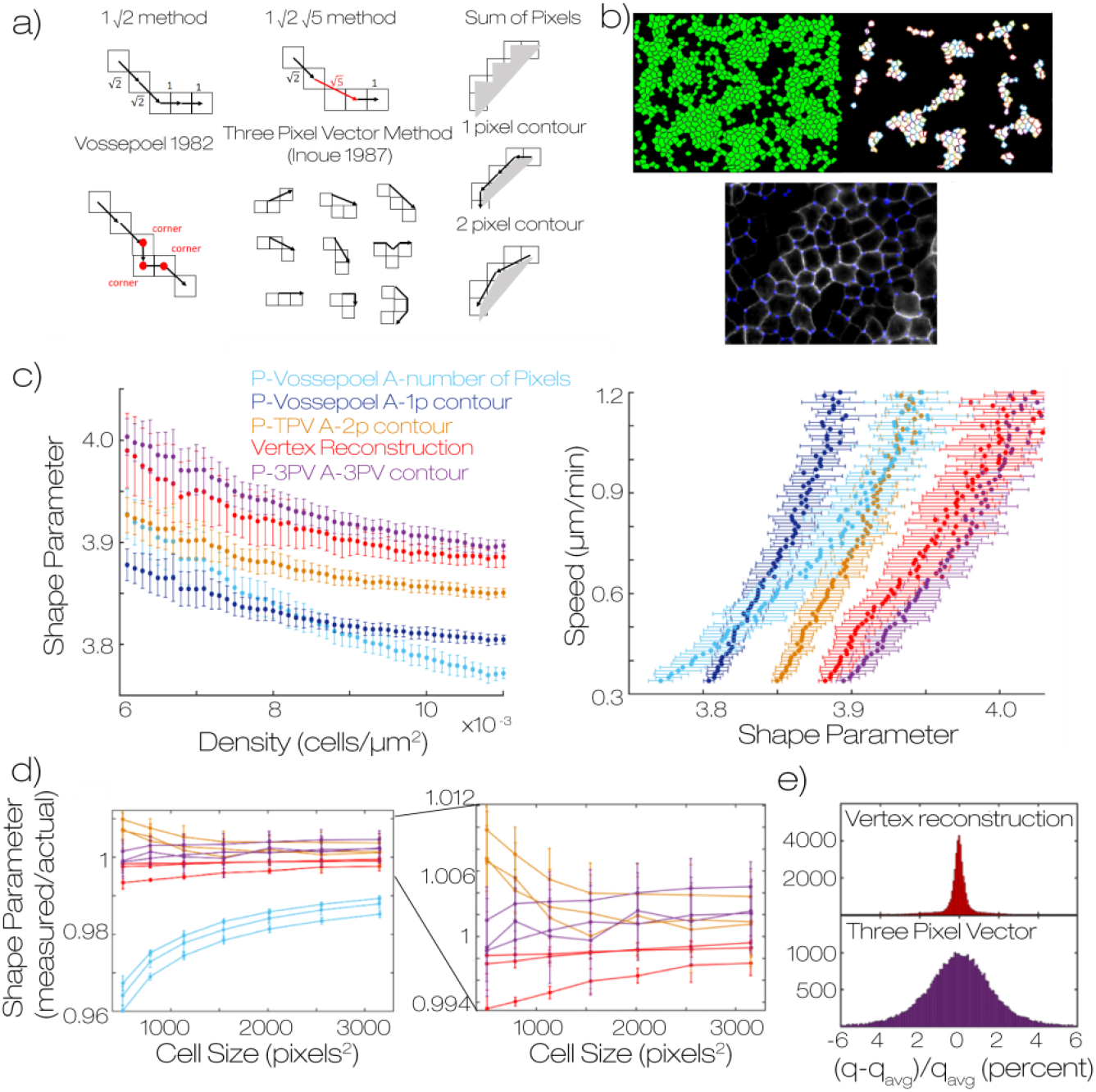
Shape metrics are resolution dependent. (a) Schematics describing different popular methods for determination of perimeter and area of a polygon constructed from pixels. Methods depicted include the 1 root 2 method – default in imageJ, and voesspoel – default in Matlab regionprops. We also schematize several methods for determining polygon area. Sum of pixels is default in both imageJ and Matlab (b) An image of cell outlines (green) with marked vertex locations (pink). The cells which do not have a complete set of neighbors are discarded and remaining polygons are constructed from the vertices. We show an overlay of detected vertices on the raw data. (c) For one experimental dataset, WT on 2mg/ml collagen – we plot shape vs density for a variety of shape metrics and Speed vs shape for the same set of metrics. Errorbar represents the standard deviation of 60 fields of view. (d) We segmented simulation data and plot shape/actual shape vs area for this data for a variety of shape metrics for images of different resolution. We observe that some shape metrics show large errors and are highly resolution dependent. We show a zoom in of the top performing metrics. Error bar represents the standard deviation of three replicates (e) We measured shape of fixed cell data and used the two top performing metrics to measure cell shape. The fixed data is a time series of images where there is no cell motion but images show photobleaching and noise variation. We know the shapes are not changing with time, however the measured shape fluctuates due to changes in noise. We measure the distribution of measurements around the mean for the two metrics for each cell. Shapes reconstructed from vertices are much more robust to noise.

**Fig. S2:**
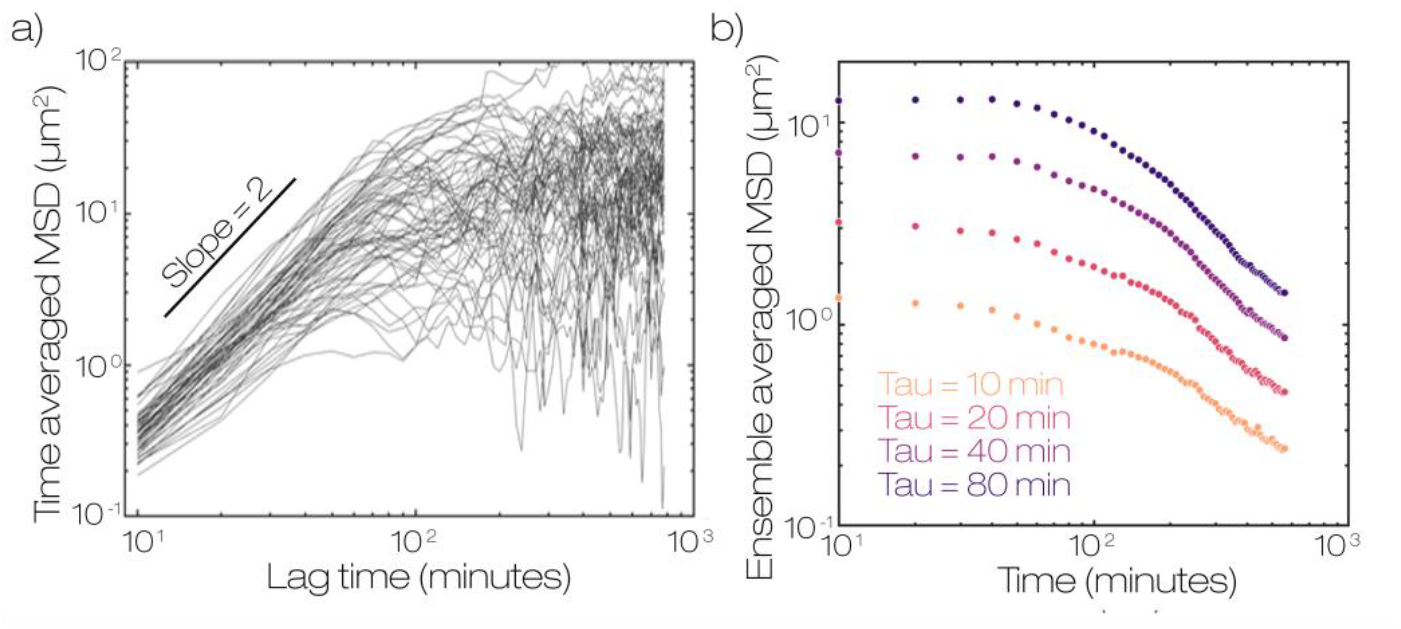
Mean square displacements are time dependent. (a) Time averaged mean squared displacements for individual cells show a plateau at long time scales indicating cells are not freely moving at long times. A small representative subset of 100 cells from one field of view is displayed (b) mean squared displacement for different lag times decay with time indicating that cell motion is reduced at later times. Data is across all cells in 60 fields of view in a single single experiment on a 2mg/ml collagen gel under wild type conditions.

**Fig. S3:**
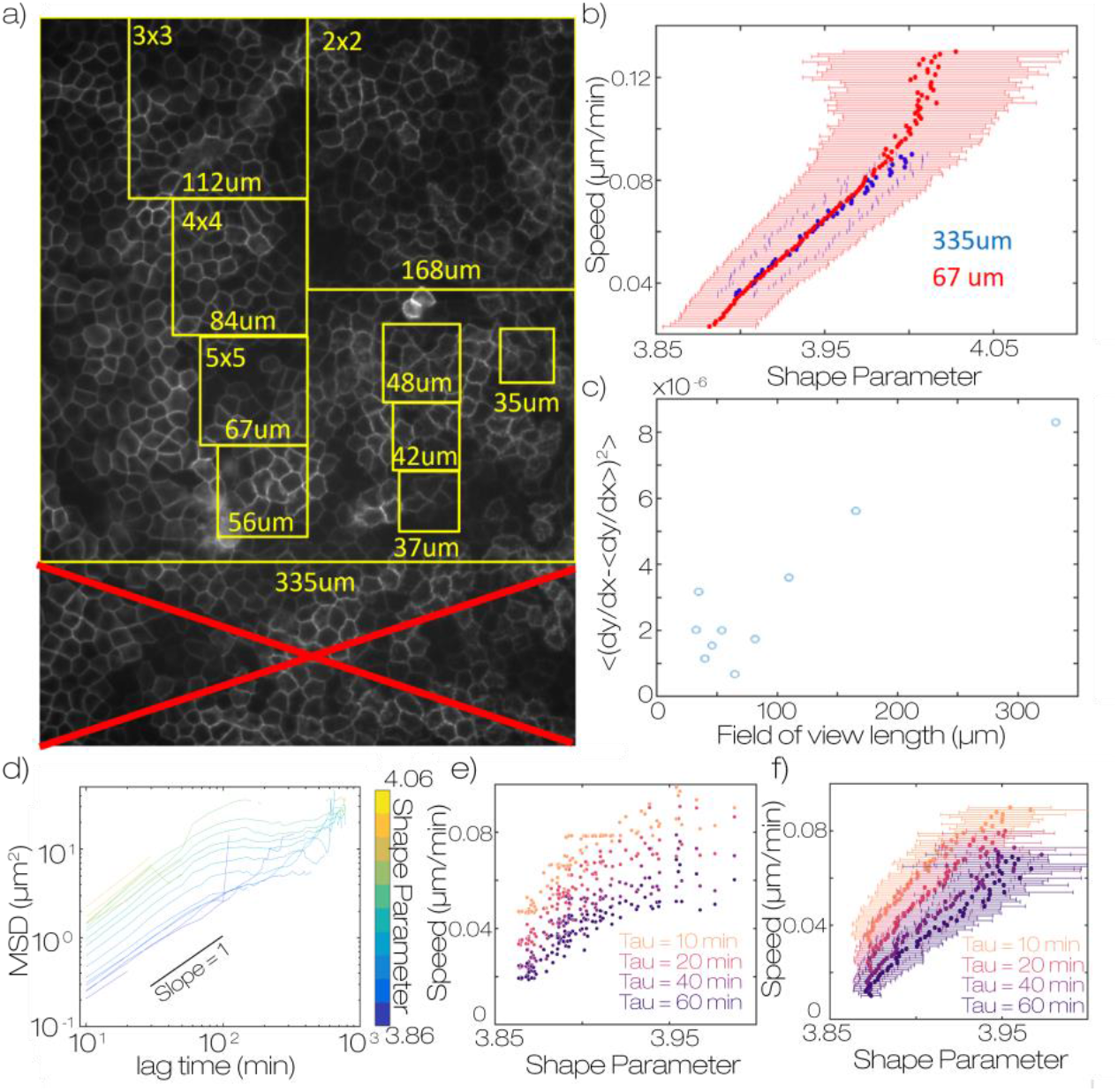
Correlation between shape and speed does not depend on the field of view size or time between images. (a) representation of different field of view sizes. For this analysis the field of view was first truncated into a square window 335×335 µm. This field was subdivided into 2×2, 3×3… 10×10 sections and each section was averaged independently as if that was the full field of view. (b) speed vs shape parameter for the full field of view 335×335 µm and a 5×5 truncation 67×67µm. Both field of view sizes show the same final average. All other truncation sizes give the same average values. Errorbars represent the standard deviation of 60 and 1500 fields respectively (c) comparison of the derivative in the linear portion of the curve in b, from q=3.93 to q = 3.97. The value is larger for large fields of view because there can be greater heterogeneity within the same field of view. At lower values the linearity plateaus due to noise from averaging fewer cells. This shows that our data is consistent with energy which is defined at the single cell level. At ∼100um or groups of a couple dozen cells we reach a noise floor. This analysis uses the TPV metric for cell shape to ensure there are enough cells within small fields of view to get a representative average. (d) Mean squared displacement for a dataset on a WT 2mg/ml collagen gel. Curves are binned according to the shape parameter at the initial time of each subtrajectory. Subtrajectories are not overlapping. Lag times are at 10 minute intervals. We observe that for any lag time there is lower displacement for lower shape parameter. (e) Shape parameter vs speed for a single field of view over 12 hours for different lag times. (f) Shape parameter vs speed averaged across all fields of view in the sample. We see that increasing the lag time shifts this curve downward consistent with the motion being diffusive. At longer times the distance traveled only increases by sqrt(t) as time increases by t leading to lower values of speed. The data appears to reach the noise floor at low shape parameter for larger tau.

**Fig. S4:**
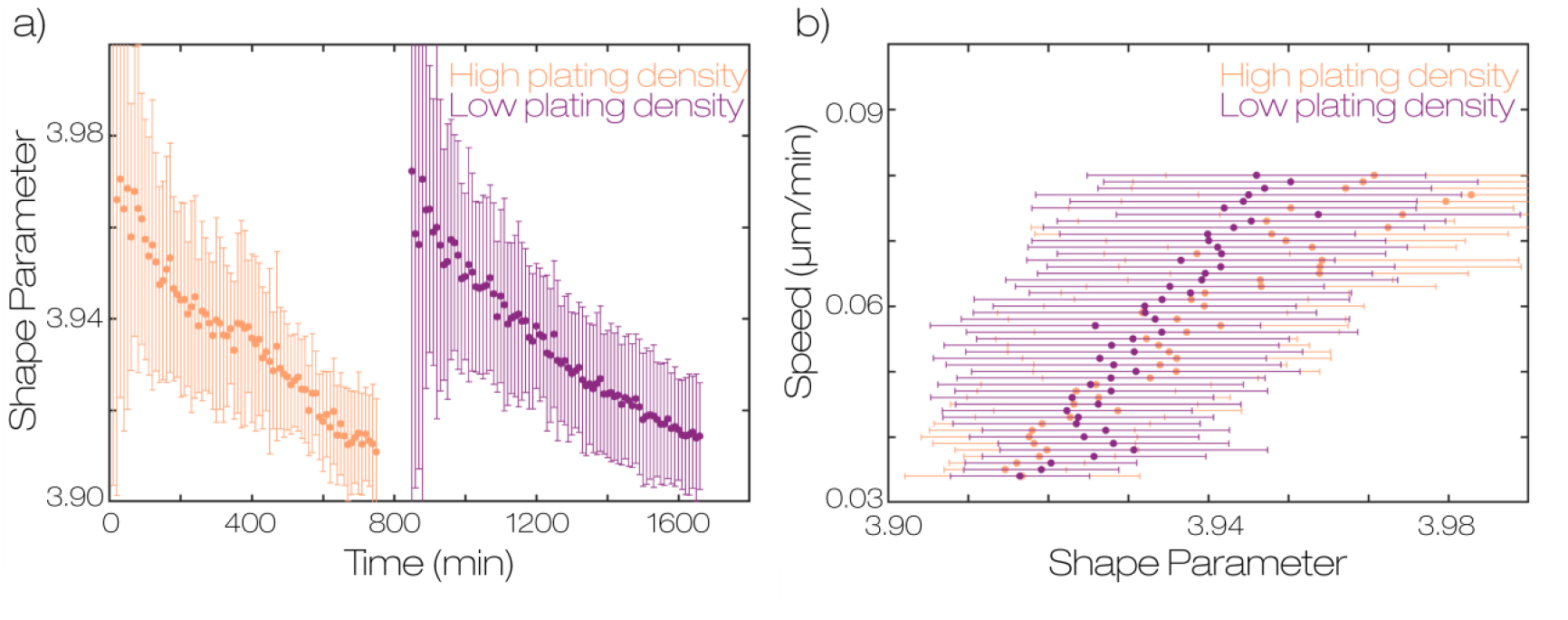
Monolayer remodeling does not depend on initial cell seeding density. (a) Cell Shape vs time for monolayers seeded at two different densities. Both samples were made at the same time with ∼ 700K cells (high density) and ∼350k cells (low density). The samples were imaged sequentially starting from the time point when the monolayer became confluent across most of the cover slip. (b) Relationship between cell shape and speed for the samples in a).

**Fig. S5:**
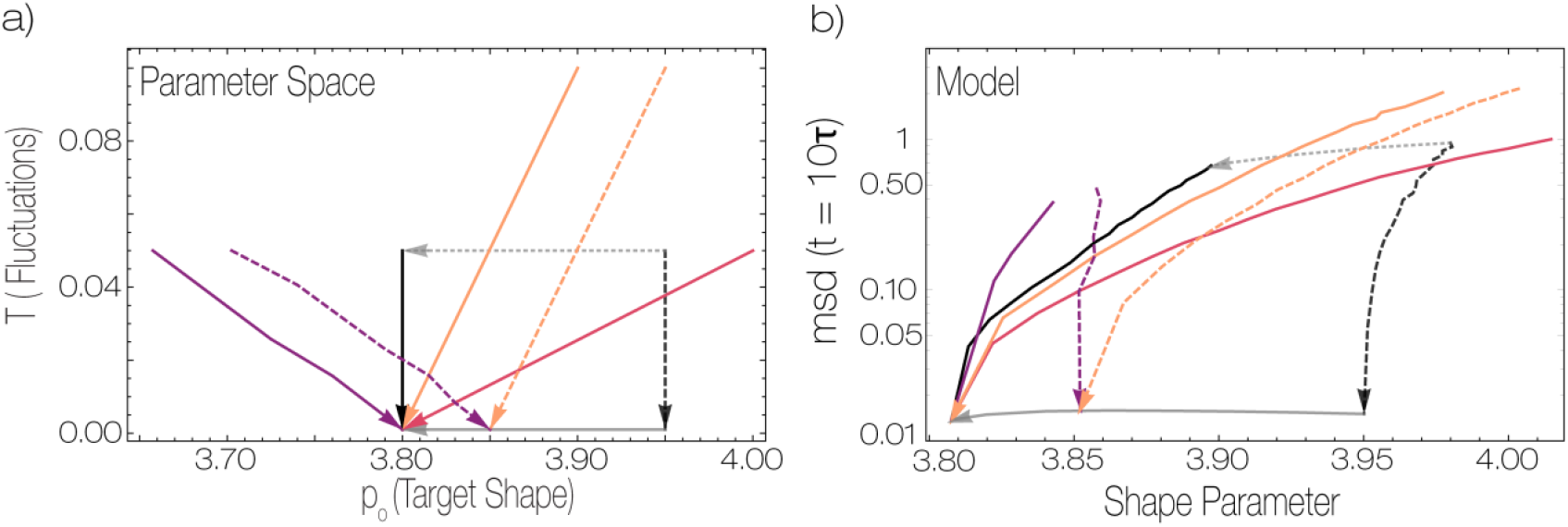
Additional modeling of relationship between cell shape and speed in Active Vertex models. (a) Parameters of the thermal Voroni model were varied along several representative curves. Along these curves a simulated monolayer was equilibrated at each point then measurements were made on the monolayer. Solid curves approach zero temperature at a value of p_0_ where the tissue is rigid, while dashed curves approach T=0 at a value of p_0_ where the tissue is floppy. Colors represent trajectories with different slopes. Dashed lines indicated different values of p_0_ approaching T=0. Arrows indicate the order of simulations along a trajectory (c) Observed values of speed (quantified by MSD in a given time window) and shape corresponding to the parameter space trajectories shown in panel b). MSD is given in units of 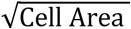 over a time window of 10 natural time units

**Fig. S6:**
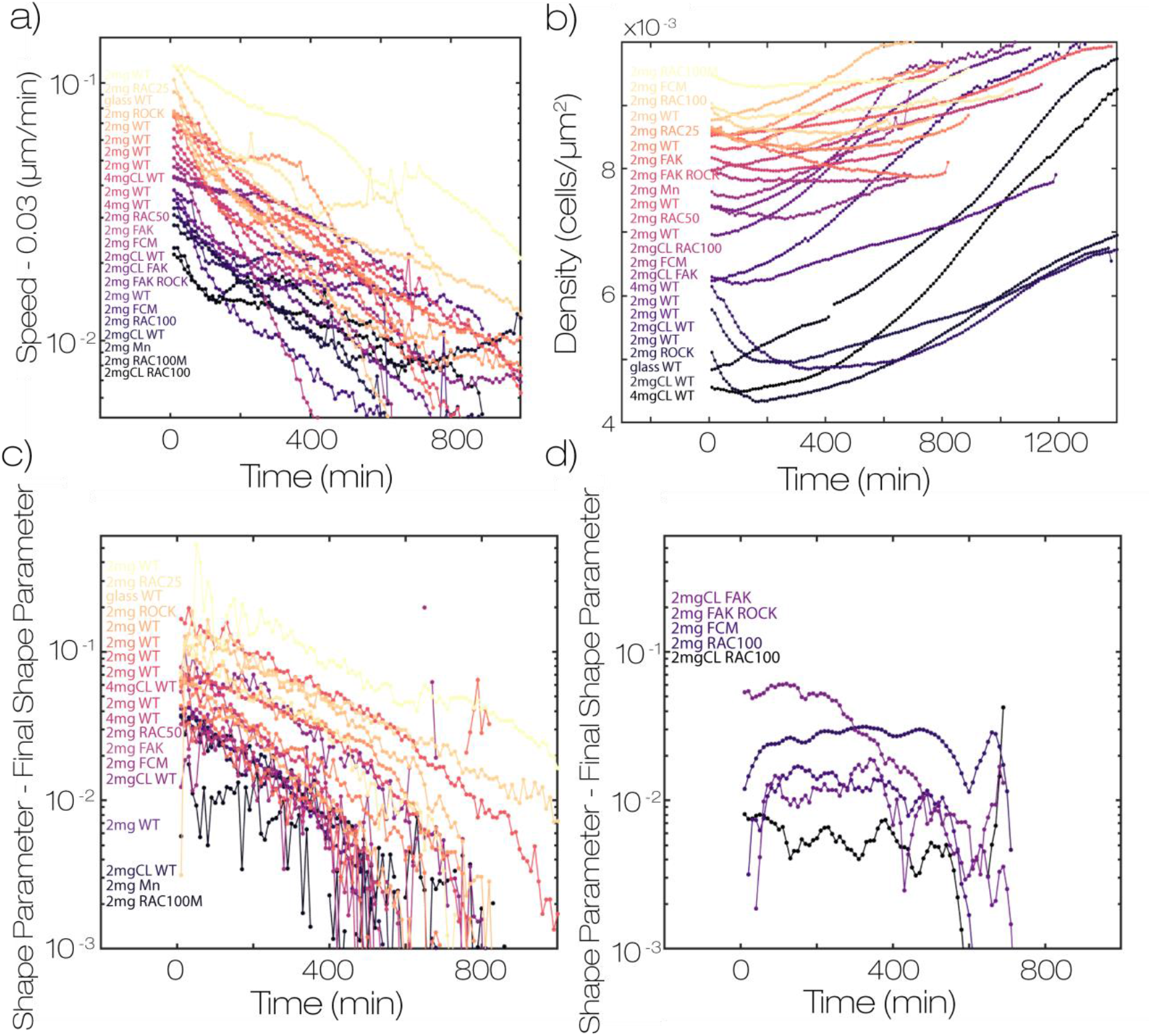
Shape, Speed and cell area decrease with time across all conditions. (a) plot of speed vs time for all conditions in figure 2b. The final speed for all the time series plateau at 0.03um/min so this value is subtracted from each time series. Curves are colored in order of speed at time = 0 min. The same color map is used in b,c (b) density vs time for all datasets. The data is colored by initial density and differs in color map from a-c. (c) plot of shape-final shape for all data. Each time series decays to the final shape with similar kinetics. (d) several datasets were too noisy and were moved to a different plot for clarity. These datasets were smoothed temporally with a 10-point window to reduce noise. These data slightly reduce in shape with time however the difference in initial and final shape are comparable to the noise in our measurement. All data points represent the time average of at least 30 fields of view in one sample.

**Fig. S7:**
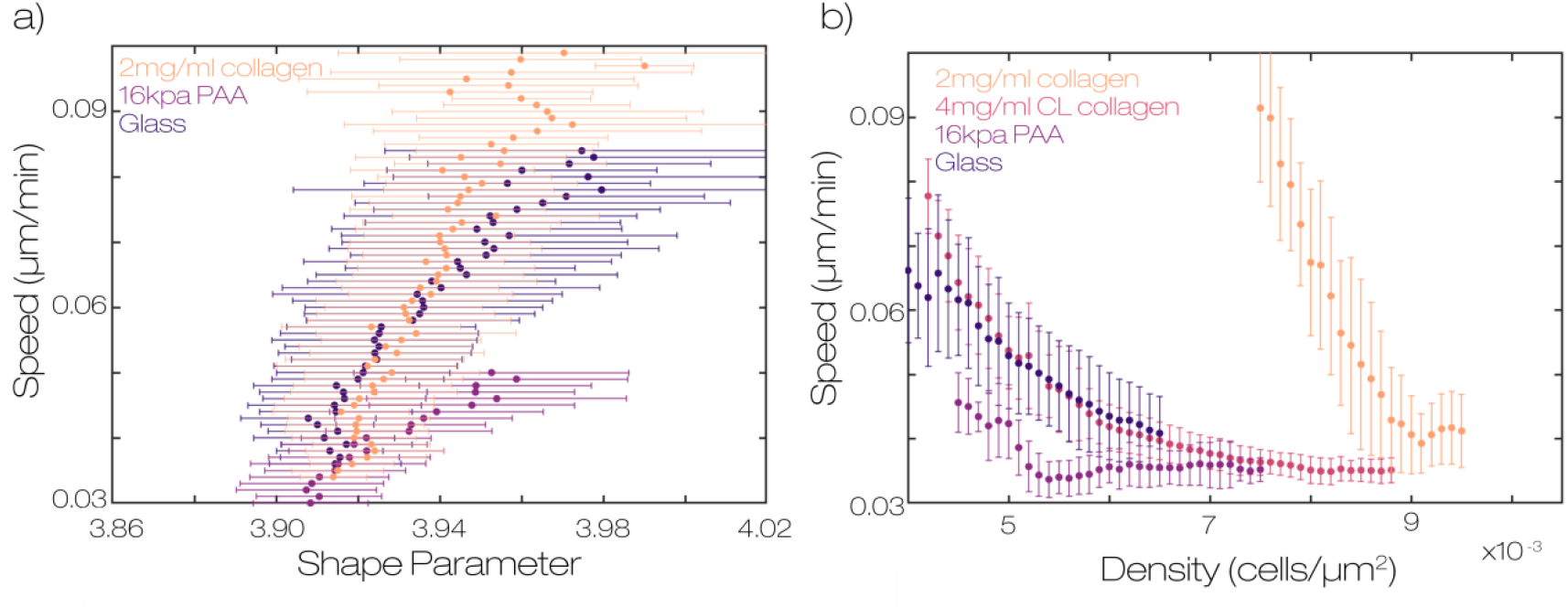
Relationship between cell shape and speed is independent of substrate stiffness. (a) relationship between shape and speed on division rate matched conditions on collagen, 16KPa polyacrylamide, and Glass (b) relationship between speed and cell density on substrates with different stiffness. Error bars represent binned averages at each speed for at least 30 fields of view over at least 60 time points.

**Fig. S8:**
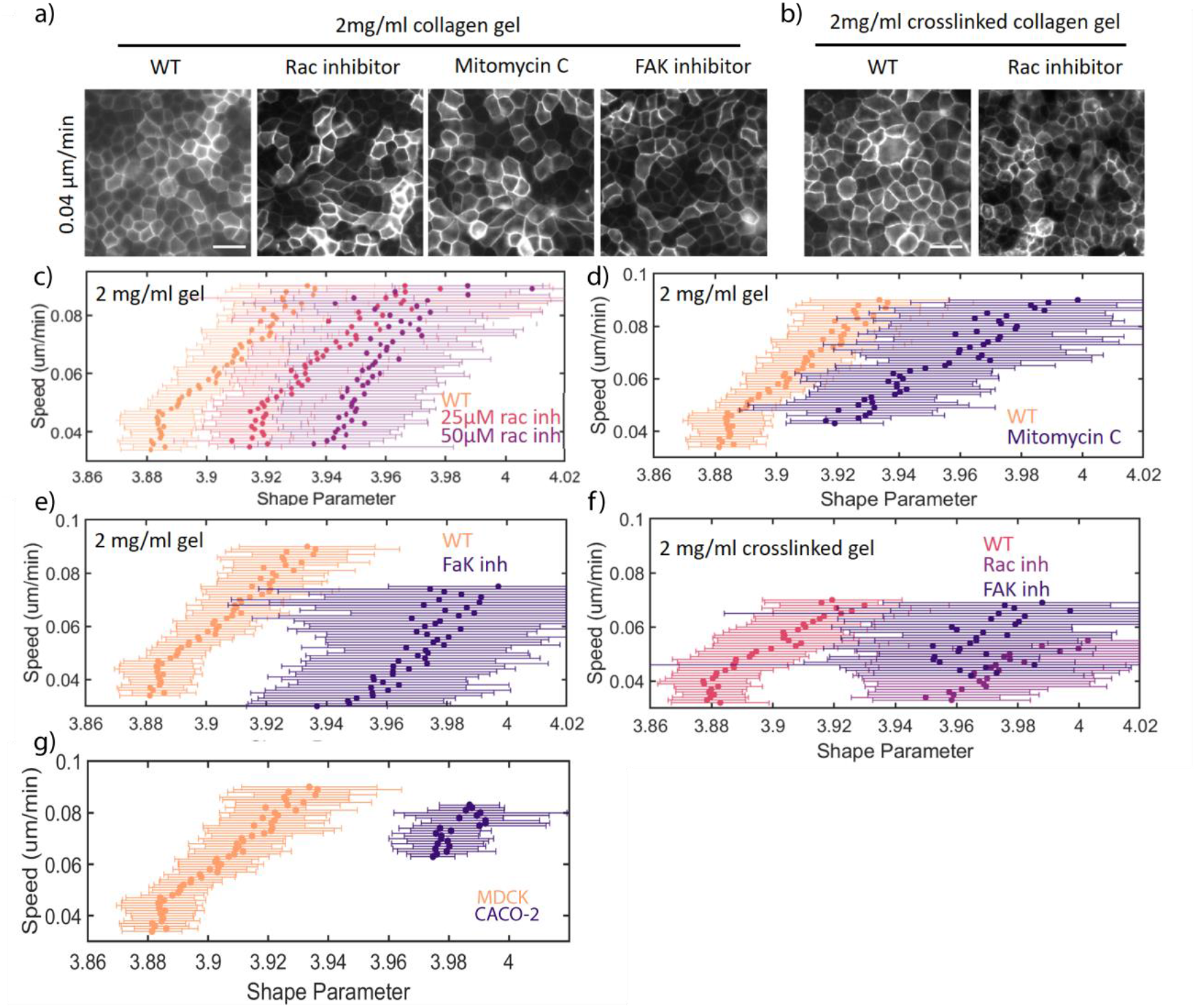
Shift in relationship between shape parameter and speed is qualitatively similar across conditions. (a) Representative images for several inhibitor conditions at low average speed. (b) Images on stiff substrates (c-e) Speed vs Shape parameter curves for different inhibitors. Similar shifting behavior is observed across these conditions. (f-g) Similar behavior is also observed on stiff substrates and for CACO-2 epithelial cells. Error bars represent binned averages at each speed for at least 30 fields of view over at least 60 time points.

**Fig. S9:**
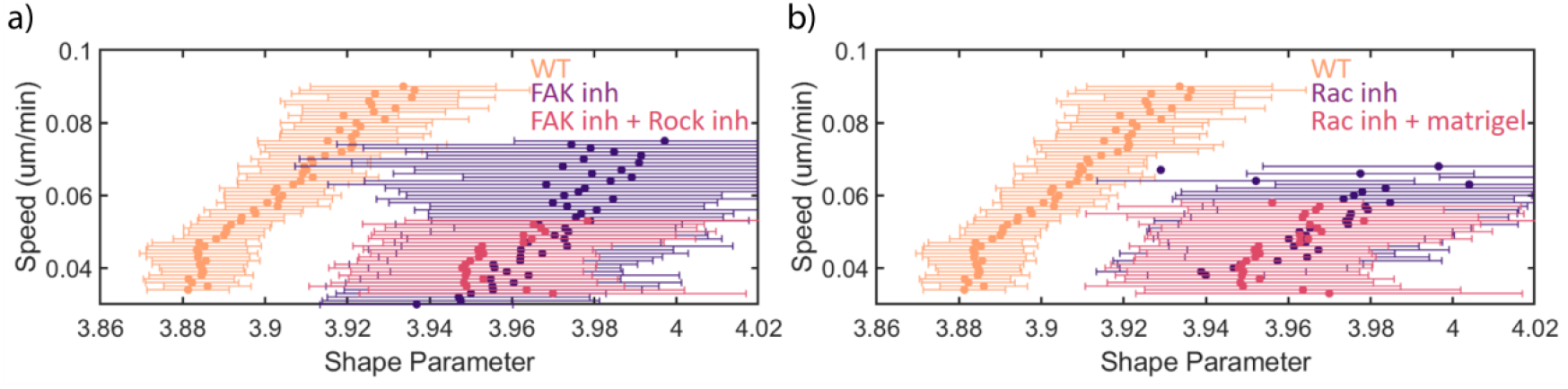
Rescue experiments do not restore shape speed correlation. (a) Comparison of shape speed correlation for cells treated with a FAK inhibitor and with the addition of a Rock inhibitor. Rock inhibition has been observed to rescue the polarity of 3D MDCK cultures with FAK knockdown. Rock inhibition has also been observed to restore collective motility in FAK knockdown cells (see discussion). (b) Comparison of shape speed correlation for cells treated with a RAC inhibitor on collagen gels and collagen gels with 1mg/ml matrigel. RAC is required for the assembly of laminin at the basal surface of 3D cultures of MDCK cells. The polarity defect of RAC knockdown cells can be rescued by providing laminin, a component of matrigel, in the substrate (see discussion). Neither experiment significantly restored the correlation between shape and speed. Error bars represent binned averages at each speed for at least 30 fields of view over at least 60 time points.

**Fig. S10:**
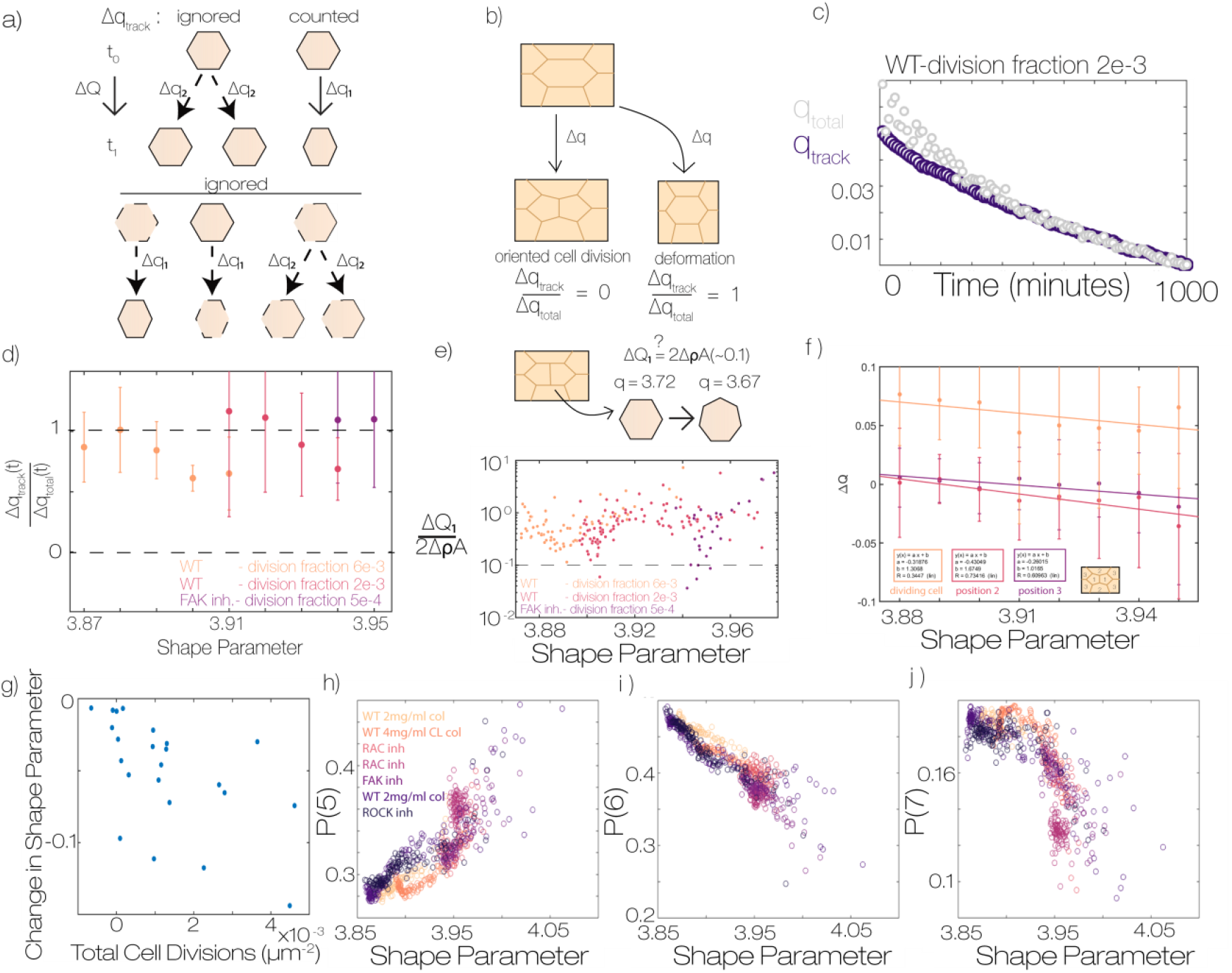
Oriented division is insufficient to explain differences in monolayer remodeling. (a) Schematic of different modes of shape change – either shape changes during a cell division Q2 or within a trajectory Q1. There are additional q1 type and q2 type shape changes which are ignored by cqq. (b) schematic of different sources of cell shape change and their relative values of q_track_ (c) for a single condition we plot q_track_ and q_total_ over time. We observe that for most of the experiment these metrics are the same. (d) ratio of q_track_ to q_total_ for three different conditions with variable division fraction noted in the legend. We observe in all cases that the ratio is close to 1 independent of cell shape. Error bars represent standard error of each bin. (e) The q_track_ metric has ignored the gain of neighbors for cells adjacent to division. We show that the value of q_track_ is several fold larger than this effect. Points represent time averaged values for at least 50 fields of view (f) average shape change per cell division from division tracking data as a function of the average cell shape. We observe larger decreases in cell shape when for the mother cell and neighbors when average cell shape parameter is large. Error bars represent standard deviation of divisions in each bin (h) plot of total cell division vs total shape change. If cell divisions cause all shape change, we would expect a strong correlation between these variables. Each point represents a different condition from Fig 5a. (g-h) relationship between shape parameter and the fraction of cells with either 5,6, or 7 neighbors for different conditions. Across these conditions we observe a common relationship between the topology of the monolayer and the shape parameter.

**Fig. S11:**
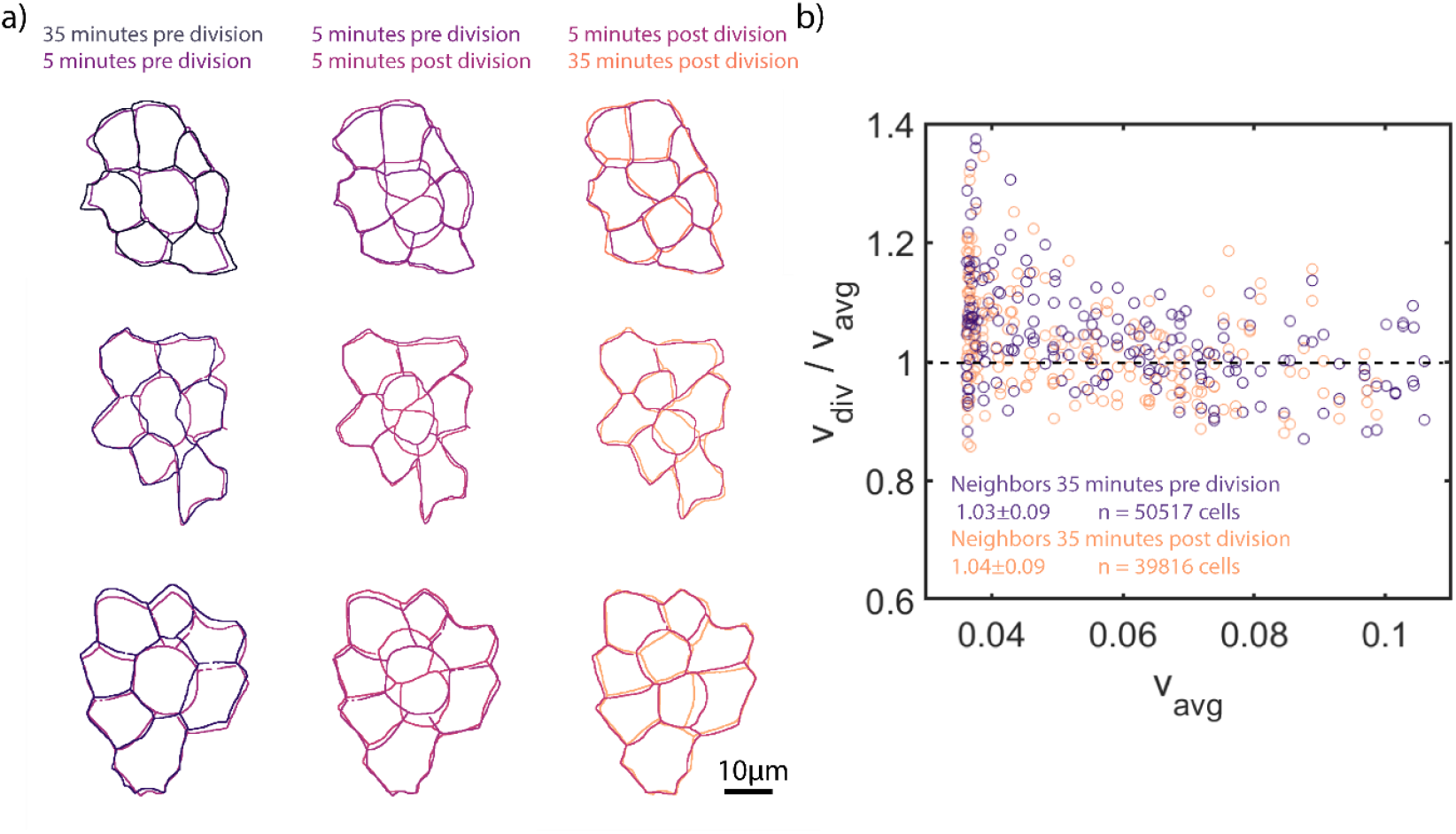
Cell divisions do not produce large deformations of the monolayer. (a) representative outlines of dividing cells in the monolayer during mitotic rounding, cytokinesis and reintegration of daughter cells into the monolayer. (b) Measurement of neighbor cell speed before and after cell divisions. All cells adjacent to a detected cell division are averaged for a given time point and compared to the average cell speed at that time point. Cell speed is the displacement of cells over a 10 minute lag time.

**Fig. S12:**
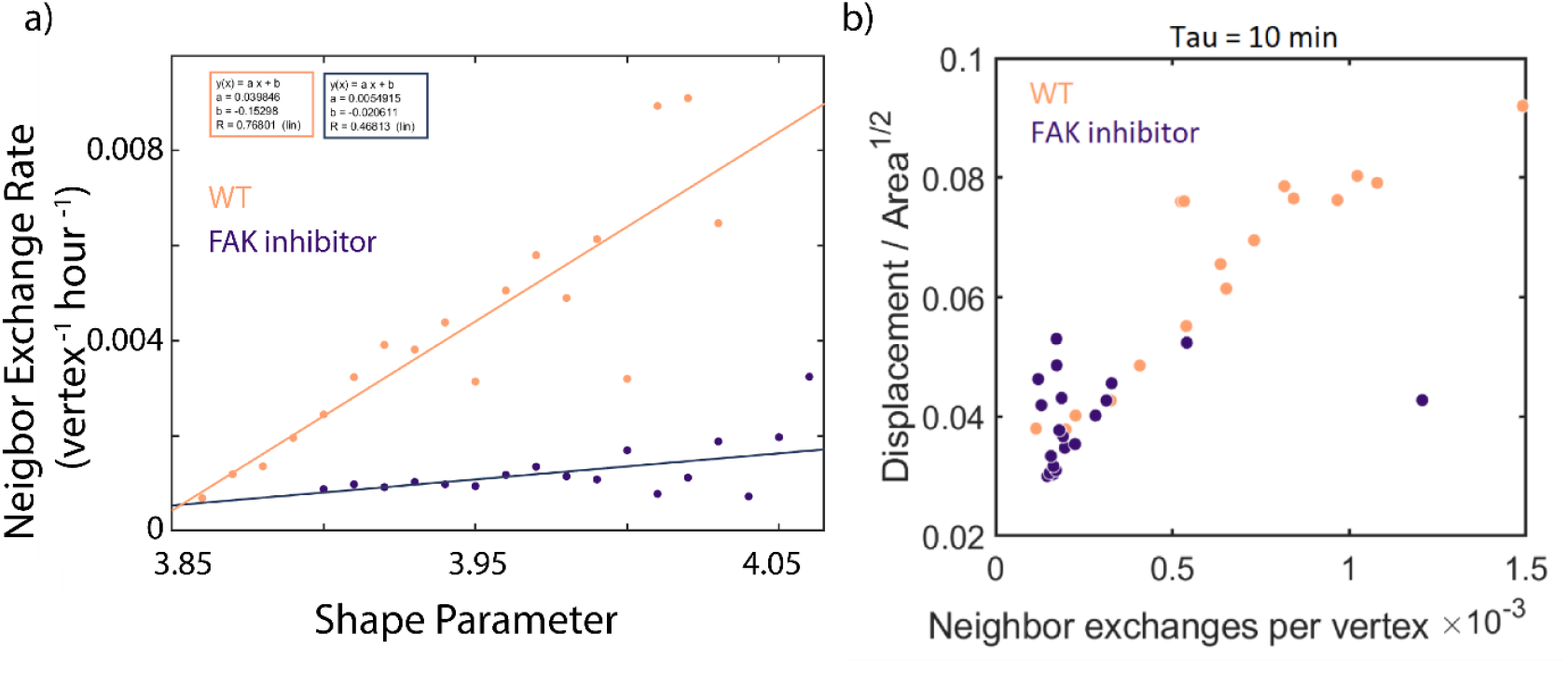
Neighbor exchange rates depend on shape and cell division rates. (a) Neighbor exchange rate vs shape for WT and FAK inhibited cells on 2mg/ml collagen gels. Neighbor exchanges were counted by identifying 4 fold vertices in the data. A 4-fold vertex represents an unstable configuration in the system and therefore will either resolve in the opposite direction it was formed (successful exchange) or back in the same direction (failed neighbor exchange). Our method does not differentiate the two types of neighbor exchange. Each point is the ratio of four fold to three fold vertices detected at the given average shape parameter across at least 50 fields of view and 80 time points in each condition. (b) relationship between neighbor exchange rates and measured cell speed for data points in a)

**Fig. S13:**
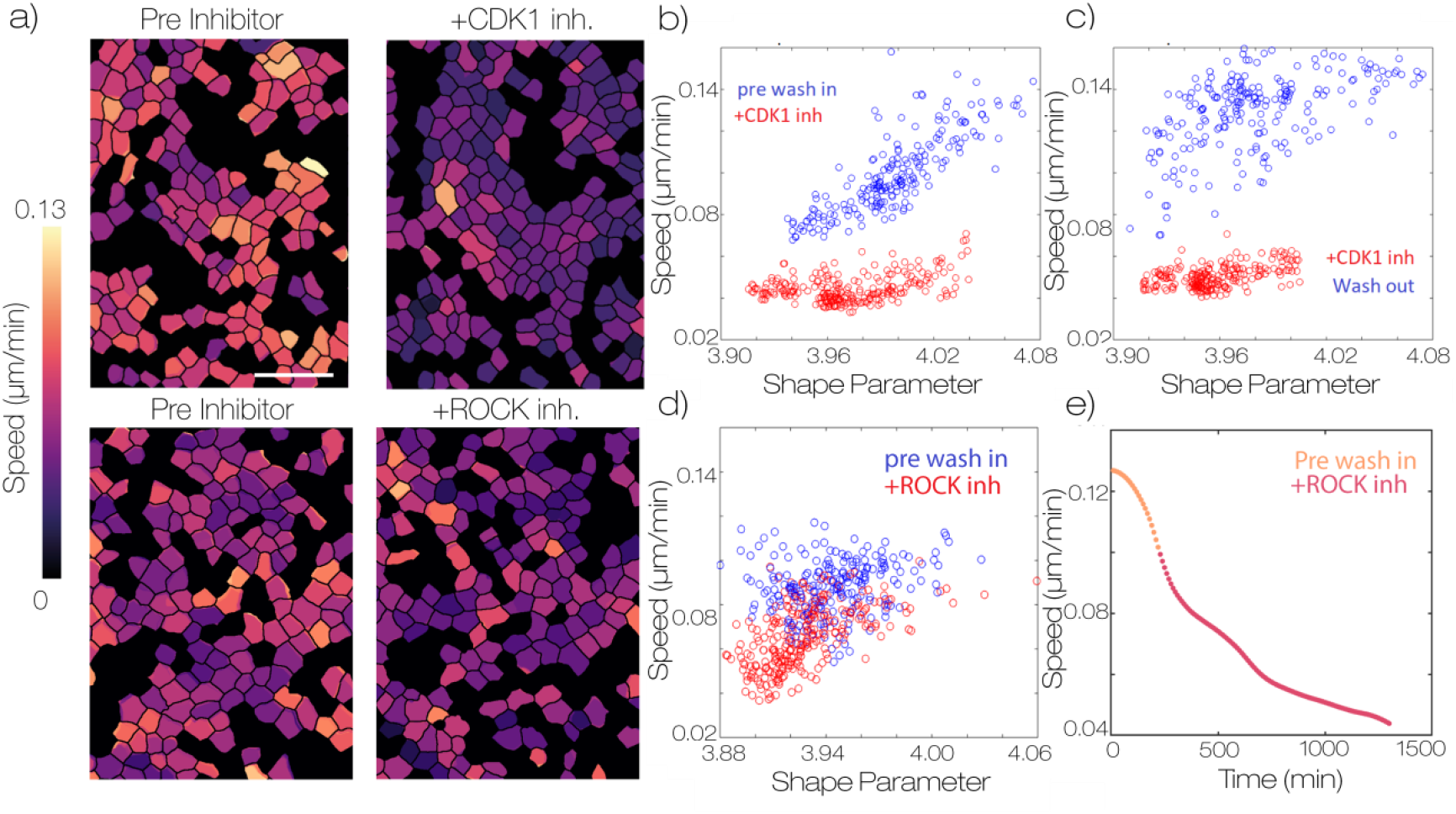
Inhibition of ROCK does not lead to large reduction in cell motility. (a) representative maps of cell speed before and after adding inhibitors of CDK1 (5µM RO-3306) and ROCK (20 µM y27632). Scale bar is 50 microns (b) measurements of cell shape vs speed for each field of view 100 minutes before and after adding the CDK1 inhibitor (c) measurements of cell shape vs speed for each field of view 100 minutes before and after washing out the CDK1 inhibitor (d) measurements of cell shape vs speed for each field of view 100 minutes before and after adding the ROCK inhibitor (e) plot of average cell speed vs time for samples treated with ROCK inhibitor.

**Fig. S14:**
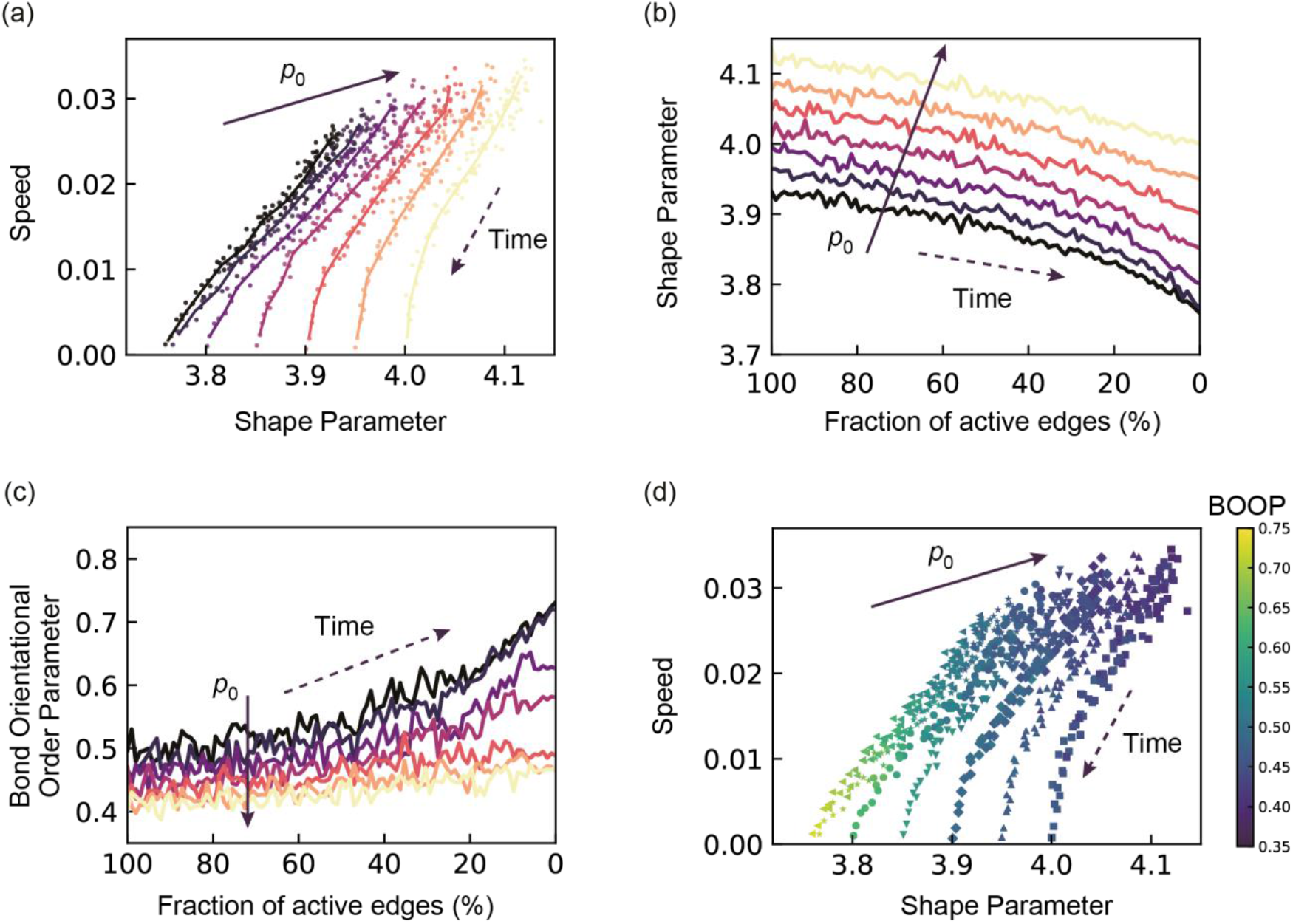
Time-evolution of the average cellular speed, the average shape parameter and the bond orientational order parameter (BOOP) for different shape indices, p_0,_ with *τ*^R^=1. (a) The average cellular speed is plotted against the average shape parameter. The curves are obtained by smoothing the data points. (b) The average shape parameter is plotted against the fraction of active edges. (c) The BOOP is plotted against the fraction of active edges. (d) The average cellular speed is plotted against the average shape parameter with the color mapping the value of the BOOP. The color is mapped as shown in the color bar. In (a-c), *p*_0_ = [3.70,3.75,3.80,3.85,3.90,3.95,4.0] from dark color to light color. In (d), the data for each *p*_0_ is respectively represented by left-pointing triangle (3.70), star (3,75), filled circle (3.80), down-pointing triangle (3.85), diamond (3.90), up-pointing triangle (3.95) and square (4.00) symbols. In (a-d), the direction of time is indicated by dashed arrows.

**Fig. S15:**
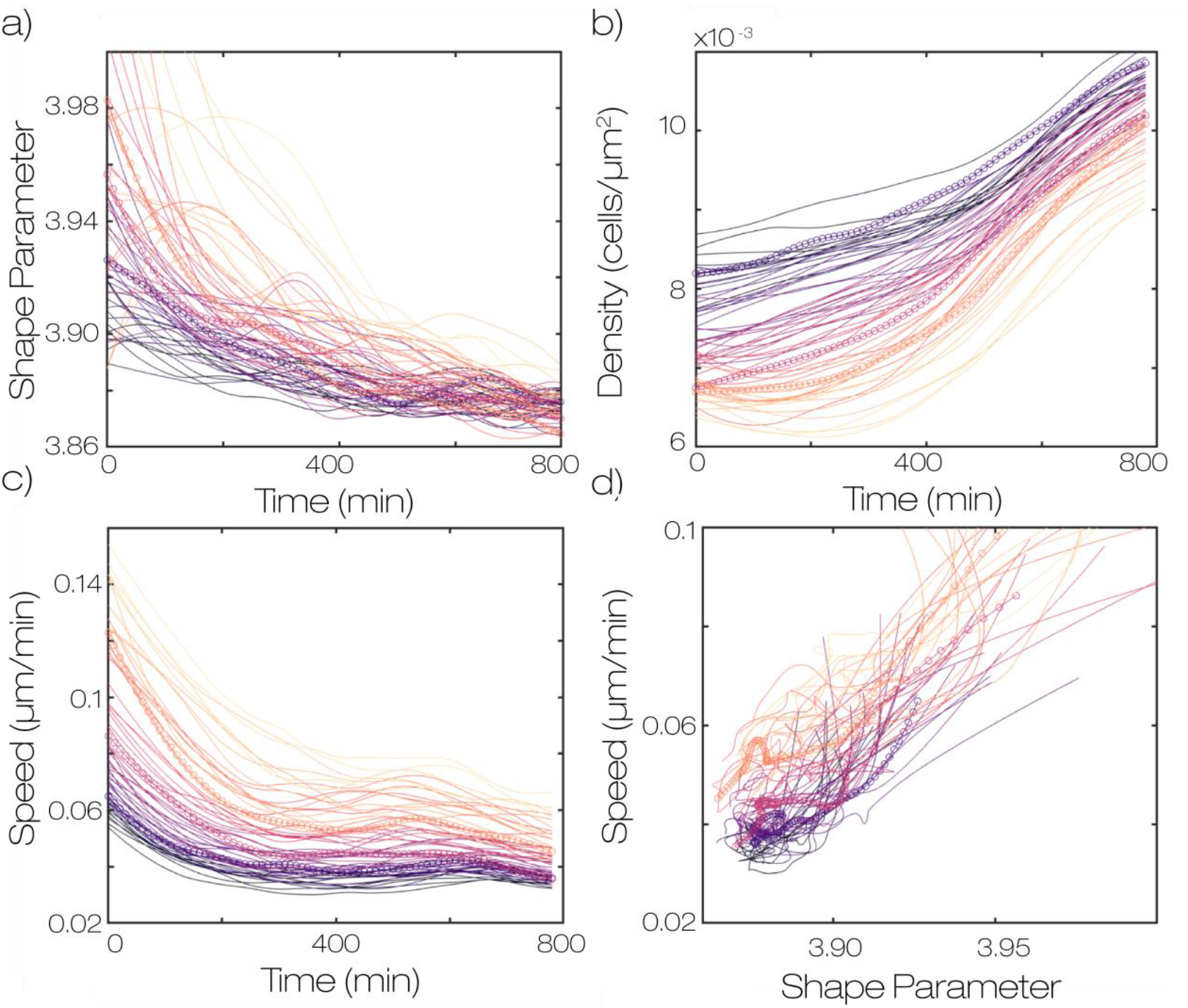
Different fields of view across the sample are qualitatively similar to the mean. (a) values of shape parameter for each field of view over time (b) density against time for the same set of fields of view. (c) speed vs time for the same set of fields of view (d) shape vs speed for the same set of fields of view. Time points are taken every 10 minutes. Data is colored according to the initial speed in the field of view. Colors are consistent across all panels. We see that the datasets which have larger initial speed also have larger initial shape and lower initial density.

**Fig. S16:**
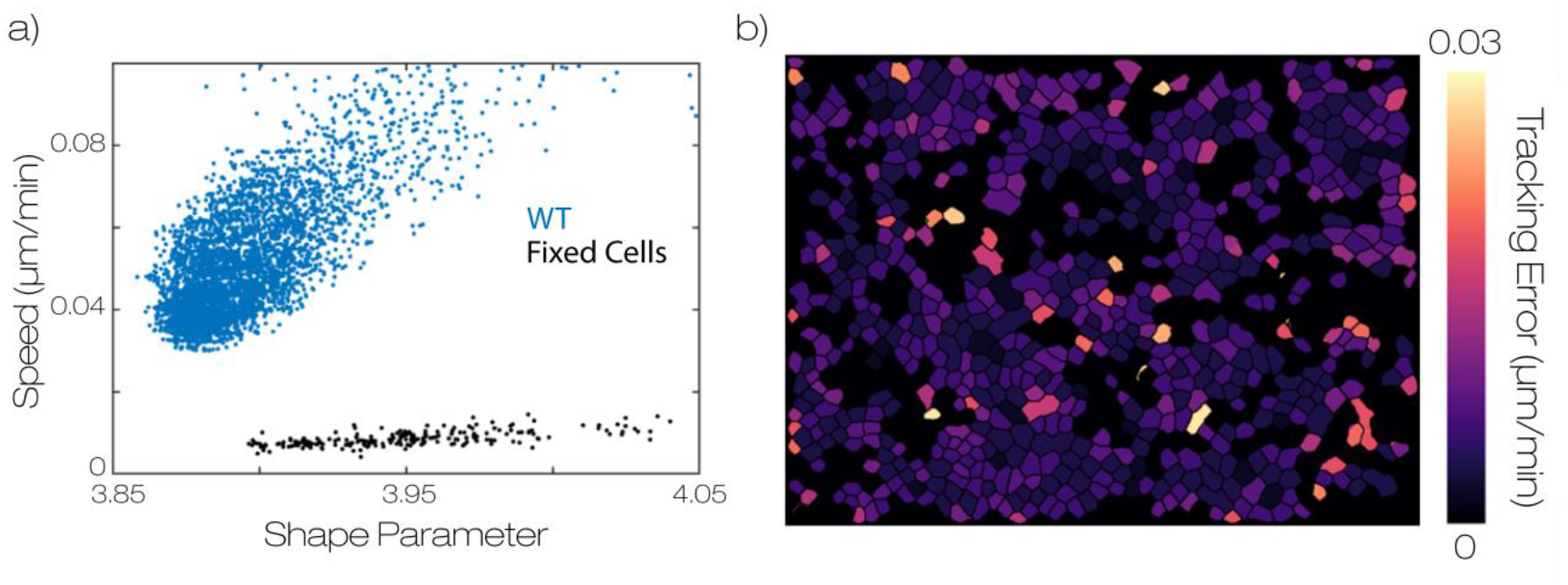
Characterization of lower bound for noise floor. (a) relationship between shape and speed in WT data on a 2mg/ml collagen gel and under the same conditions but fixed in 4% PFA for 10 minutes before imaging. We observe an estimated noise floor with minor shape dependence and average value well below the speeds measured in the experiment. The noise floor may be larger due to larger fluctuations in protein levels and cell heights in live samples. Points represent the average cell shape and speed at a single field of view and time point (b) heat map of perceived displacements in fixed data. Measurement error is fairly homogeneous and very few single cells tracking errors comparable to experimental displacements.

## Legends for Movies S1 to S8

**Movie S1:** Representative time series from a WT monolayer on a 2mg/ml collagen gel. Tracked displacements and cell shapes are plotted as colormaps on the outline generated from segmentation. Time step is 10 minutes.

**Movie S2:** Representative time series from a WT monolayer on a 2mg/ml collagen gel (left) and on a 4mg/ml crosslinked gel (right). Time step is 10 minutes.

**Movie S3:** Representative time series from a monolayer treated with 100 µM RAC inhibitor NSC23766 on a 2mg/ml collagen gel. Tracked displacements and cell shapes are plotted as colormaps on the outline generated from segmentation. Time step is 10 minutes.

**Movie S4:** Representative time series from a WT condition (left) and with 50:50 Fibroblast conditioned medium to normal culture medium (right). Time step is 10 minutes.

**Movie S5:** Representative time series showing wash in and wash out of the CDK1 inhibitor RO-3306. The inhibitor was added at 5µM at 5:00 and washed out of the imaging chamber at 10:00. Colormap of cell speed is plotted on the segmented cell outlines. Time step is 10 minutes.

**Movie S6:** Representative time series showing behavior after addition of microtubule destabilizing compound Nocodazole. Nocodazole was added at 300nM at 4:00. Time step is 10 minutes.

**Movie S7:** Representative time series showing behavior after addition of DNA crosslinker Mitomycin C. Mitomycin C was added at 20µg/ml at 7:30. Time step is 10 minutes.

**Movie S8:** Representative movie showing the segmentation of a time series. Segmented cells are represented in pink and overlaid on the GFP signal. Time steps is 10 minutes. Movie duration is 800 minutes.

